# Histones are exosome membrane proteins regulated by cell stress

**DOI:** 10.1101/2024.04.08.588575

**Authors:** Birendra Singh, Marcus Fredriksson Sundbom, Uma Muthukrishnan, Balasubramanian Natarajan, Stephanie Stransky, André Görgens, Joel Z. Nordin, Oscar P. B. Wiklander, Linda Sandblad, Simone Sidoli, Samir EL Andaloussi, Michael Haney, Jonathan D. Gilthorpe

**Author notes:** Contributed equally and should be considered as first authors (Birendra Singh, Marcus Fredriksson Sundbom). Considered as co-senior authors (Uma Muthukrishnan, Balasubramanian Natarajan).

## Abstract

Histones are conserved nuclear proteins that function as part of the nucleosome in the regulation of chromatin structure and gene expression. Interestingly, extracellular histones populate biofluids from healthy individuals and when elevated may contribute to various acute and chronic diseases. It is generally assumed that most extracellular histones exist as nucleosomes, as components of extracellular chromatin. We analysed cell culture models under normal and stressed conditions to identify pathways of histone secretion. We report that core and linker histones localize to extracellular vesicles (EVs) and are secreted via the multivesicular body/exosome pathway. Upregulation of histone EV secretion occurs in response to cellular stress, with enhanced vesicle secretion and a shift towards a population of smaller EVs. Most histones were membrane associated with the outer surface of EVs. Degradation of EV-DNA did not impact significantly on EV-histone association. Individual histones or histone octamers bound strongly to liposomes and EVs, but nucleosomes did not, showing histones do not require DNA for EV binding. EV histones colocalized most frequently with the tetraspanin CD63 but using genetic or pharmacological intervention, we found that all known pathways of exosome biogenesis acted positively on histone secretion. Inhibition of autophagy and lysosomal degradation had a strong positive effect on EV histone release. Unexpectedly, EV-associated histones lacked the extensive post-translational modification of their nuclear counterparts, suggesting loss of PTMs may be involved in their trafficking or secretion. Our data does not support a significant role for EV-histones existing as nucleosomes. We show for the first time that histones are secreted from cells as membrane proteins via EVs/exosomes. This fundamental discovery provides support for further investigation of the biological activity of exosome associated histones and their role in disease.

## INTRODUCTION

Histones are basic proteins^1^ abundant in the eukaryotic nucleus. The core histones (H2A, H2B, H3 and H4) associate with DNA to form the nucleosome^2, 3^ and together with linker histone H1 they form chromatosomes^4^. Hence, the overwhelming bulk of investigation into the localization and function of histone proteins has focused on their roles in chromatin organization and gene regulation within the nuclear compartment. Functions for cytoplasmic and extracellular histones have also been described^5, 6^. Perhaps the best understood paradigm for histone release by activated cells is in association with modified chromatin during neutrophil extracellular trap (NET) formation, by a specific pathway of regulated cell death known as ‘NETosis’^7–11^. However, our understanding of pathways for histone secretion in non-activated cells is lacking.

Elevated levels of extracellular histones are found in biofluids from critically ill patients with conditions such as acute lung injury/acute respiratory distress syndrome (ALI)/(ARDS), sepsis, trauma, severe COVID-19 infection and mediate inflammation through toll-like receptor (TLR)-4 and TLR-2 signaling^12–19^. Increased circulating histones have been implicated in poor survival as well as in organ dysfunction^13, 20–22^, mechanical ventilation, secondary infection^23^, thromboembolic events and acute kidney injury^13, 14, 20–24^. The toxic effects of extracellular histones can be meliorated, for example by degradation using activated protein C, or blocking using anti-histone monoclonal antibodies or C1 esterase inhibitors. These approaches have indicated therapeutic potential, with increased survival and reduced organ dysfunction^24, 25^. However, relatively low levels of circulating histones are also found in biofluids under normal physiological conditions^26–29^. This raises important question of how histones reach the extracellular environment and whether histones are normally secreted by healthy cells, as well as whether there is a function for extracellular/circulating histones in the non-disease state.

Histones are frequently identified as components of extracellular vesicle (EV) proteomes^30–32^, including microvesicles/ectosomes or apoptotic vesicles (generated via outward budding of the plasma membrane), apoptotic bodies (released via membrane blebbing), amphisomes^33^, and exosomes^30, 34, 35^. Exosomes are endosome-derived vesicles^36^ formed as intraluminal vesicles from inward budding within multivesicular bodies (MVBs) and released from cells upon the fusion of an MVB with the plasma membrane^37^. They carry numerous proteins^34^, bioactive lipids^38^ RNAs^39^ and DNA^40^. Two basic pathways for exosome biogenesis have been defined; an endosomal sorting complex required for transport (ESCRT)-dependent and an ESCRT-independent pathway requiring ceramide^41, 42^. It is unclear whether extracellular histones are associated with a particular type of EV, whether they localize to the surface or within EVs, or how they are trafficked to a secretory pathway.

The presence of histones in purified EVs has been suggested to be due to contamination with apoptotic vesicles that co-purify^34^, or fusion of EVs with free histones during ultracentrifugation^33^. Another suggestion has been that histones are sorted into the lumen of EVs with exosome DNA (exoDNA) in a nucleosome-like configuration^31, 43^, with micronuclei into exosomes^44^, or are secreted via mitochondrial derived EVs^6^. Hence, the dominant viewpoint is that EV associated histones are nucleosomes, or chromatin associated particles that bind to EVs^30, 31^. Whether this is the case for all EV histones, or all cell types, remains an important question.

We have previously identified histones as secretory components of injured rat brain tissue^45^. Significant progress has been made in understanding pathways of EV/exosome biogenesis using rodent brain oligodendrocyte cell lines^46–48^. Oligodendrocyte EVs contain both core and linker histones^47^ and can be grown readily in 2-dimensional (2D) or 3D cell culture systems under serum free conditions to facilitate the purification a sufficient quantity of EVs for the biochemical characterization of cargo proteins, even for histones where there is a significant risk of contamination with abundant cellular proteins such as histones^49^. Hence, we used this as a starting point to define whether and where histones localize to EVs and to define how they are secreted.

In this study, we used the rat oligodendrocyte progenitor cell line OLN-93^50^ as a model to elucidate the pathways for EV-mediated histone secretion. Using a combination of experimental strategies, we show for the first time that core and linker histones are present in MVBs and released via exosomes. Histone secretion was upregulated in response to different stress paradigms. With stress there was a shift to a population of EVs of smaller diameter and higher floating density. Surprisingly, most EV histones were strongly associated with the EV membrane and interacted directly with membrane lipids in liposomes. In accordance with a previous study^30^, the majority EV-DNA associated with the EV surface. However, we found that EV-DNA and EV-histones could be degraded independently. Both ESCRT-dependent and –independent, as well as autophagosome-lysosomal pathways, were involved in the routing of histones for secretion. Using imaging flow cytometry, we could show both H3 and H4 colocalize to the surface of tetraspanin positive EVs, predominantly a CD63 positive population. Intriguingly, secreted EV associated histones lacked post-translational modifications, distinguishing them from their nuclear, chromatin associated, counterparts.

## MATERIALS AND METHODS

All materials and reagents were obtained from Thermo Fisher Scientific or Merck/Sigma-Aldrich unless otherwise stated.

### Cell lines and culture

Cells were maintained in a humidified incubator at 37°C/5% (v/v) CO_2_. OLN-93^50^, HEK/HEK293T (ATCC CRL-3216/CRL-1573TM) and NSC-34 (CEDARLANE) cells were cultured in high glucose Dulbecco’s modified Eagle’s medium (DMEM). HeLa (ATCC CCL-2) and Caco-2 (ATCC HTB-37) cells were maintained in in Eagle’s Minimum Essential Media, AC16 (ATCC CRL-3568) in DMEM:F12 and A549 (ATCC CCL-185) in F12K. Complete media were routinely supplemented with 2 mM L-glutamine, 10% (v/v) heat inactivated FBS with 100 IU/ml penicillin/streptomycin.

An OLN-93^EGFP-CD^^63^ stable cell line (EGFP fused to the intraluminal N-terminus of human CD63)^51^ was established by selection in 1 mg/ml G418 sulphate^52^. A construct for mouse H3.1 (Hist1h3a, P68433) was generated by RT-PCR from mouse cDNA and cloned (details available on request) into a modified doxycycline-inducible lentiviral vector pCW57.1 (a gift from David Root, Addgene plasmid # 41393). A HEK293–H3.1 stable cell line was generated via transduction of lentiviral particles and selection with 2.5 μg/ml puromycin (Sigma) for 4 days. Histone expression was induced by addition of 1 μg/ml doxycycline (Sigma) for 24 h.

### Antibodies

Detailes of the antibodies used in this study are enlisted in Supplementary table S1.

### Western blot

Protein concentrations were estimated using BCA Protein Assay Kit (Pierce/Thermo Fisher Scientific) and Qubit (Thermo Fisher Scientific) proteins were separated by SDS-PAGE on AnykD Criterion™ TGX™ precast gels (Bio-Rad) and transferred to 0.2 µm nitrocellulose membranes by using a Trans-blot Turbo instrument (Bio-Rad). Blots were blocked with 2.5% BSA in PBS and then incubated with primary antibodies as indicated above, prior to washing (3x PBS containing 0.001% v/v Tween-20) and detection with HRP-conjugated secondary antibodies (Vector Labs or Dako), followed by detection with ECL Prime Western Blot Detection (GE Healthcare). Blots were imaged on an Odyssey®CLx system (LI-COR) or ChemiDoc XRS+ (Biorad).

### EV production and isolation

For 2D culture and EV production in flasks under normal and stress conditions, cells were seeded in tissue culture flasks coated with 50 µg/ml poly-D-lysine and grown to confluence in complete media. Cells were washed 3x with warm PBS and growth media was replaced with serum-free Opti-MEM media, which was conditioned for 24 h. Where necessary, EVs were concentrated from a large volume of conditioned media by tangential cross flow diafiltration (100 kDa MWCO, Sartorius).

For 3D culture and EV production at scale under control conditions only, OLN-93 cells grown in a hollow fibre bioreactor (HFBR; FiberCell Systems, USA). Cells were seeded/maintained and harvested according to the manufacturer’s protocol. Complete DMEM was used as a reservoir feeding media. The cell compartment of the HFBR was supplemented with OptiMEM supplemented with 10% (v/v) Exosome-Depleted FBS (Gibco/Thermo Fisher Scientific). EV harvesting was performed at a regular 12 h interval and overgrowth of the cells was controlled by flushing the cell compartment, as described in the manufacturer’s instructions. EVs produced using the HFBR were qualitatively indistinguishable from EVs produced in cell culture flasks.

EVs were isolated from conditioned medium first cleared by sequential centrifugation^53^ at 500 xg for 10 min, 2,000 xg for 20 min, 10,000 xg for 30 min (to remove cell debris and larger microvesicles) followed by a 0.2 µm filtration step. EVs were pelleted by ultracentrifugation at 100,000-110,000 xg for 90 min (Optima XPN-100 ultracentrifuge using a SW28 or SW32Ti rotor, Beckman Coulter) at 4°C. EVs were subject to 2 washes in 50 mM 4-(2-hydroxyethyl)-1-piperazineethanesulfonic acid (HEPES)/150 mM NaCl (pH 7.4) with recovery at 100,000 xg for 90 min. EVs (referred to as EV^100,000^ ^xg^) were used directly after resuspension, or stored in the same buffer at -80°C after snap freezing in liquid nitrogen.

Protein or DNA content were quantified with a Qubit 3.0 fluorometer using the appropriate Qubit protein, or dsDNA HS assay kit (Life Technologies/Thermo Fisher Scientific).

### OptiPrep equilibrium density gradient centrifugation

Iodixanol (OptiPrep) density gradients were used to fractionate EV^100,000^ ^xg^. In brief, a 12-36% (w/v) OptiPrep gradient was prepared in 25 mM HEPES (pH 7.4), 150 mM NaCl, 0.25 M sucrose, 1 mM EDTA and 10 mM Tris-HCl (pH 7.4). EV^100,000^ ^xg^ (300-400 μg based on protein content) were placed in the bottom 36% layer of a 6-step gradient in a 13.2 ml open top polyallomer tube (331372, Beckman Coulter). Following ultracentrifugation at 100,000 xg for 18 h/4°C, (SW41 Ti rotor), fractions were collected, and density was estimated by comparison to a reference gradient using the absorbance at 340 nm. OptiPrep was removed by washing with 25 mM HEPES (pH 7.4), 150 mM NaCl for 4 hours at 100,000 xg. EV pellets were resuspended in PBS or HEPES buffer. In some cases, fractions 4-7 were pooled and in this case are referred to as called EV^OptiPrep^.

### Immunofluorescence staining and Duolink Proximity Ligation Assay (PLA)

OLN-93 or HEK293T were grown in complete media on poly-D-lysine coated glass coverslips and switched into Opti-MEM for 16 h prior to fixation and staining. Formaldehyde was added to a final concentration of 4% (w/v) and incubated for 15 min at room temperature (RT). Cells were then washed three times with PBS and blocked in 10% FBS/0.1% (v/v) Triton-X 100 for 1 h at RT. Primary antibodies were added in blocking solution and incubated overnight at 4°C. Samples were washed 3x 5 min in PBS/0.1% (v/v) Triton-X 100 followed by addition of Alexa fluor conjugated secondary antibodies. After 1 hour at RT, coverslips were again washed 3x with and then mounted on glass microscope slides with Dako Fluorescence Mounting Medium and imaged using a confocal microscope (Nikon Eclipse C1).

For PLA, permeabilization, blocking and primary antibody incubation was carried out as described above. DuoLink PLA (Sigma Aldrich) was performed according to the manufacturer’s protocol. In brief, PLA probes were incubated with the samples for 1h at 37°C in a humidified chamber, followed by 30 min ligation and 100 min amplification. After washing, samples were incubated with DAPI and briefly washed in ddH2O. Cover slips were mounted and imaged as described above.

### Cellular stress

Cells were seeded in tissue culture flasks (T-25 or T-175) or on coverslips coated with poly-D-lysine and grown to confluence in complete DMEM. Cells were then washed 3x with warm PBS and switched to serum-free conditions in Opti-MEM. For heat stress experiments shown in Fig. 2A and S10, the conditioned medium was collected before heat stress (Control/Before sample; 24 h conditioning period) before subjecting the cells to heat stress at 42°C. After a 3 h heat shock, the cells were returned to 37°C for a 4 h recovery period. EV-conditioned medium was collected (Heat stress sample; 7 h conditioning period) and again replaced with fresh Opti-MEM. Conditioned medium was then collected after a further 17 h (After sample; 17 h conditioning period).

For other stress experiments, media was changed before exposure to the stressor and control samples were maintained in parallel with the treated samples. Media was harvested after a 24 h conditioning period): Heat stress (42°C for 3 h), lipopolysaccharide (LPS) stress (1 µg/ml LPS, *E. Coli* O111:B4 for 24 h), hypoxia (humidified 2% (v/v) O_2_, 5% (v/v) CO_2_, balance N_2_) for 24 h though O_2_ equilibration occurs slowly in liquid media due to the slow absorption of O_2_ at a static interface water^54^, oxidative stress (single application of 50 µM H_2_O_2_) which provides a transient oxidative stress due to the rapid degradation of H_2_O_2_ in cell culture media^55^.

### Cell viability assays

OLN-93 cells were seeded on to coverslips coated with poly-D-lysine and grown to 80-90% confluency. Cells were washed 3x with warm PBS and switched to Opti-MEM. Following exposure to stress conditions, media was removed, and cells were stained with Hoechst 333421 (5 µg/ml) and propidium iodide (50 µg/ml) in PBS for 20 min at RT. Cells were washed 1x with PBS and coverslips were mounted and imaged by confocal microscopy (Nikon Eclipse C1).

Apoptosis was quantified using the Incucyte S3 Live Cell Analysis platform (Sartorius). OLN-93 cells were seeded (7000 cells/well) in 24-well plates (Corning #3526) coated with poly-D-lysine. Cells were incubated overnight then washed 3x with PBS and switched to Opti-MEM. Annexin V-red and Cytox-green reagents (Sartorius) were added according to the manufacturer’s recommendations. The cells were subjected to stress conditions and then imaged under phase contrast and green/red fluorescence channels every 2 h over a 24 h period. The number of single and doubled labelled cells were quantified in each channel and using IncuCyte Base software (Sartorius).

### StrataClean pull down of proteins

Cell culture media was cleared by sequential 500, 2000, and 10,000 xg centrifugation steps followed by 0.2 µm filtration. StrataClean resin (Agilent) was added to the cleared media (5 µl/ml). Thereafter, samples were placed on a roller for 15 h at 4 °C. The resin was collected by centrifugation 5,000 xg for 10 min. Total proteins bound to the resin was resuspended in 1X SDS-PAGE loading buffer containing 2 mM DTT, heated to 95 °C for 10 min and separated by SDS-PAGE, prior to WB.

### Nuclear and cytoplasmic fractionation

OLN-93 cells were pelleted by centrifugation and resuspended in 10 mM HEPES (pH 7.9), 1.5 mM MgCl_2_, 10 mM KCl, 2 mM DTT with Protease Inhibitor Cocktail (Roche Diagnostics) and incubated for 10 min on ice. Cells were pelleted at 2000 xg at 4°C and resuspended in the same buffer, followed by homogenization with a Dounce homogenizer. After centrifugation at 10,000 xg for 10 min at 4°C the cleared supernatant was collected as the cytoplasmic extract and the pellet was resuspended in 20 mM HEPES (pH 7.9), 1.5 mM MgCl_2_, 500 mM NaCl, 25% (v/v) glycerol, 0.5 mM EDTA, 1 mM DTT and protease inhibitors and incubated for 30 min at 4°C with continuous mixing. The mixture was then centrifuged, and the supernatant collected as the nuclear extract.

### Histone extraction

Acid extraction of histones was performed with the addition of 0.4 M H_2_SO_4_ for 2 h at RT^56^. Following centrifugation, trichloroacetic acid (TCA) was added to the supernatant to a final concentration of 20% (w/v). The mixture was incubated for 2 h at RT before centrifugation at 16,000 xg and 4 °C for 30 min. The pellet was retrieved and washed 3x with acetone before air drying. Histones were dissolved in ddH_2_O.

### Nanoparticle tracking analysis

Nanoparticle tracking analysis (NTA) was used to determine the size distribution and number of particles^57^ using a NanoSight NS300 instrument (Malvern Panalytical) equipped with NTA 3.1 software. For all our recordings, a camera level of 8-9 and a detection threshold of 3-4 were used. Samples were diluted in ddH_2_O or buffer to achieve a particle concentration of approximately 2x10^8^-1x10^9^ particles/ml. Five 60 s films were analysed for each sample and measurements were repeated for 3-5 replicate samples.

### NaCl or lithium 3,5-diiodosalicylate treatment of EVs

NaCl^58^ and Li-diiodosalicylate^59^ were used for extraction of proteins from purified EVs (10 µg in 20 mM Tris-HCl, pH 7.5) using NaCl (250 mM-750 mM) or Li-diiodosalicylate (5-50 mM). The final volume was adjusted to 25 µl using Tris-HCl. Samples were incubated for 12 h at 4°C with gentle shaking. The next day, 5 µl of the sample was taken for TEM analysis and 20 µl was subjected to ultracentrifugation at 150,000 xg in a TLA-100 rotor and benchtop ultracentrifuge (Optima Max-XP, Beckman Coulter) for 1h at 4°C. Supernatant and pellet (resuspended in 20 µl Tris-HCl buffer) were quantified for DNA (dsDNA assay, Qubit), or histone H3 by SDS-PAGE and WB. NaCl treated supernatants and pellet samples were also analysed by dot blot, where 20 µl of supernatant/pellet samples were mixed with 30 µl of Tris-HCl buffer (pH 7.4) and blotted on to a 0.2 µm nitrocellulose membrane (Biorad) using 96-well vacuum manifold (Biorad). After blotting, dot blots were treated as for WB membranes for protein detection.

### Nuclease or protease treatment of EVs

DNase I, S1 nuclease, and Exonuclease III (Thermo Fisher Scientific) were added to EV^OptiPrep^(10 µg; 2 U of DNase I, 100 U of S1 nuclease, and 20 U of Exonuclease III) in combination. For combined enzyme digestion, S1 nuclease and Exonuclease III buffers were both used at 1x concentration. Digestion mix was incubated at 37°C for 2 h. The enzymatic reaction was terminated by addition of 1x RIPA, 5 mM DTT, and boiled for 10 min at 95°C. and samples loaded on a SDS-PAGE. The concentration of histone H3 was estimated by WB. For DNA quantification, firstly nucleases were deactivated by addition of 10 mM EDTA and heating at 65 °C for 10 min. Total DNA was extracted and purified by CeleCTEV tumour DNA enrichment kit (Hansa Biomed). DNA extracted from EVs before and after nuclease digestion was also analysed by TapeStation 4200 (Agilent), using reagents and screed Tape D5000 according to the manufacturer’s instructions.

For trypsin digestion, EV^100,000^ ^xg^ from OLN-93 or HEK293-H3 were suspended in 50 mM HEPES buffer (pH 7.5) and treated for 30 min. at 37°C with Trypsin Protease (MS Grade, Pierce/Thermo Fisher Scientific) at ratios of 1:1500, 1:500 and 1:150 µg trypsin:µg protein (determined in pilot experiments to span a suitable range for partial to complete digestion of histones) in a total volume of 60 µl. Control EVs were incubated without trypsin. Trypsin was inhibited using 10 µM phenylmethylsulfonyl fluoride (PMSF, Roche Diagnostics) in a total volume of 1ml PBS for 10 min on ice. EVs were washed in PBS and pelleted at 110,000 g for 1 h prior to WB.

### Lipid isolation and liposome preparation

Lipids from intact OLN-93 cells were extracted as described by Bittame *et al.*^60^. In brief, cells were trypsinized from 5x T-175 culture flasks grown to 90 % confluency and collected by centrifugation. Pellets were then resuspended in 3 ml of 1:2 chloroform:methanol and incubated for 30 min with vortexing every 5 min. Chloroform (500 µL) and 900 µL deionized water were added and samples were centrifuged at 1300 xg then incubated for 10 min at RT. The organic bottom phase was collected reextracted with 500 µl chloroform and 900 µl deionized water, after which the sample was centrifuged again at 1300 xg for 10 min at RT. The organic phase was collected and dried with a gentle stream of nitrogen under constant rotation. Samples were dried further under vacuum for 3 h. Dried lipids were rehydrated overnight in 50 mM HEPES (pH 7.4), 150 mM NaCl and sonicated briefly in a sonicating water bath. Following rehydration, liposomes were homogenized using a 100 nm membrane mini extruder (Avanti Polar Lipids). The sample was heated to 65°C and then passed through a 100 nm membrane 30 times. Liposome size distribution and morphology were confirmed by NTA and TEM, respectively.

### Liposome/EV binding assays

Human recombinant histones from New England Biolabs: H1 (M2501S), H2A (M2502S), H2B (M2505S), H3.1 (M2503S) and H4 (M2504S), total histones extracted from calf thymus (H9250, Type IIA Sigma-Aldrich), or recombinant His-tagged octamer (BPS-52037) or nucleosomes (BPS-52038) were purchased from BPS/Nordic Biosite. For binding studies,10 µg of liposomes (in 50 mM HEPES (pH 7.4),150 mM NaCl,) were mixed with 1 µg histones and incubated for 1 h at RT with gentle mixing. Histones without liposomes was used as controls. Samples were loaded onto 100 kDa MWCO centrifugal filters with PES membrane (Amicon Ultra, Millipore) and washed 7x with buffer by centrifugation. Samples retained by the filter were analysed by WB. Liposome interaction with histones, nucleosomes, or octamers was differentiated on sucrose gradients. Histones (5 µg) were mixed with 10 µg liposomes. Nucleosomes and octamers (5 µg) were mixed with 20 µg liposomes. Total reaction was performed as a 100 µl volume in a 50 mM HEPES (pH 7.4) containing150 mM NaCl. Liposome-histone/octamer/nucleosome samples were loaded at the bottom of a 40% sucrose layer, with a discontinuous 6-step gradient up to 0% sucrose. Samples were centrifuged at 100,000 xg for 16 h/4°C using a TLS-55 swinging-bucket rotor and benchtop ultracentrifuge (Optima Max-XP, Beckman Coulter). Fractions were collected and loaded directly on SDS-PAGE prior to WB.

For dot blotting, EV^OptiPrep^ were diluted to 50 µg/ml in PBS. EVs were diluted in a 2-fold series in a 96-well microtiter plate in triplicate. Solutions were then blotted on to a nitrocellulose membrane using a 96-well vacuum manifold. After blotting, membranes were blocked in PBS containing 2.5% (w/v) BSA, followed by overnight incubation of blots in 1 µg/ml octamer/nucleosome in the PBS containing 2.5% BSA. Nucleosome/octamer binding were detected by anti-His tag and HRP conjugated anti-rabbit secondary antibodies.

### Genetic or pharmacological intervention in cell lines

All silencer RNA (siRNA) including Tsg101 (196580) Alix (s221953), Smpd2 (189939) and negative control siRNA; 4390771 were purchased from Thermo Fisher Scientific. OLN-93 cells were grown to 90% confluency on poly-D-lysine coated 25 cm^2^ flask. Cells were washed with PBS and switched to OptiMEM. A total of 300 pmol siRNA was mixed with 6 µL Lipofectamine 3000 in Opti MEM media and incubated at room temperature for 15 min prior to addition to cells. The siRNA-lipofectamine mix was incubated with cells at 37°C for 6 h and then replaced with fresh OptiMEM and further incubated for 24 h before harvest/analysis. Conditioned media was processed by StrataClean pull as described above.

For pharmacological intervention, drug concentrations were selected from previously published protocols (Supplementary table S2). Cell viability was evaluated in OLN-93 cells grown in 96-well plates treated with in poly-D-lysine using the MTT assay following treatment with a range of drug concentrations, applied in OptiMEM to a set of 6 replicate wells. After 24 h of incubation, 10 µl freshly prepared MTT (5 mg/ml in ddH_2_O) was added to each well. Cells were incubated for 3 h at 37°C. Media was aspirated and 50 µl DMSO was added to each well to dissolve formazan crystals with shaking at 200 r.p.m. Absorbance at 560 nm was read in a plate reader (Biotech Synergy 2). Wells without cells were used as controls.

Analysis of histone secretion was analysed in OLN-93 cells grown in poly-D-lysine coated T-25 flasks (80-90% confluency). Cells were washed with PBS and supplemented with 2.5 ml OptiMEM per well containing compound or vehicle (control): 10 mM 3-methyl adenine (3-MA, Sigma), 5 µM neutral sphingomyelinase inhibitor GW4869 (SelleckChem), 200 mM trehalose (Merck), 50 µM imipramine (SelleckChem), 25 µM rapamycin (SelleckChem), 100 nM Bafilomycin A1 (SelleckChem), 15 µM Dyngo (SelleckChem), 10 mM valproic acid (Sigma-Aldrich), 10 mM sodium butyrate (Sigma-Aldrich), 200 nM Trichostatin A (Sigma Sigma-Aldrich), 20 nM romidepsin (SelleckChem), 50 µM YF-2 (SelleckChem), 15 µM C646 (SelleckChem), 10 µM A-485 (SelleckChem). After 24 h treatment, supernatant (conditioned media) was removed from the flasks. Conditioned media (2 mL) was taken and processed by centrifugation at 500 xg, 2,000 xg and10,000 xg followed by protein pull down using StrataClean resin.

### High resolution single EV imaging flow cytometry

EVs were analysed at a single vesicle level by high resolution imaging flow cytometry (IFCM; Amnis Cellstream, Cytek; equipped with 405, 488, 561 and 642 nm lasers) based on previously optimized settings and protocols^61–63^. Briefly, fluorescently conjugated anti-histone and anti-tetraspanin antibodies were used to probe for respective proteins on the EV surface in pre-cleared conditioned media (CM) samples collected from HeLa cells for control and H_2_O_2_ treatment. For each measurement, 25 µl of CM (and unused media as a control) were incubated overnight at room temperature with the antibodies listed above. For detergent lysis controls, samples were treated with 0.5% (v/v) NP-40 for 30 min at RT post staining. All samples were diluted 50-500-fold in PBS-HAT^64^ before acquisition via the autosampler of the Cellstream instrument with FSC turned off, SSC laser set to 40%, and all other lasers set to 100% of the maximum power setting. Samples and controls were acquired for 3-5 minutes at a flow rate of 3.66 µl/minute with CellStream software version 1.2.3 and analysed with FlowJo Software version 10.8.1 (FlowJo LLC). Fluorescence calibration and reporting of fluorescence data in molecules of equivalent soluble fluorophores (MESF) was performed as described before^61, 62^. Dulbecco’s PBS pH 7.4 (Gibco) was used as instrument sheath fluid.

### Histone analysis by mass spectrometry

Histones were prepared as described previously by Shechter *et al*.^56^. Briefly, histones were acid-extracted with chilled 0.2 M sulfuric acid (5:1, sulfuric acid:pellet) and incubated with constant rotation for 4 h at 4°C, followed by precipitation with 33% TCA overnight at 4 °C. The pellet was dissolved in 50 mM ammonium bicarbonate (pH 8.0), and histones were subjected to derivatization using 5 µl of propionic anhydride and 14 µl of ammonium hydroxide to balance the pH at 8.0. The mixture was incubated for 15 min and the procedure was repeated. Histones were then digested with 1 µg of sequencing grade trypsin (Promega) diluted in 50 mM ammonium bicarbonate (1:20, enzyme:sample) overnight at room temperature. The derivatization reaction was repeated to derivatize peptide N-termini. The samples were dried in a vacuum centrifuge.

Samples were resuspended in 10 µl of 0.1% trifluoroacetic acid and loaded onto a Dionex RSLC Ultimate 300 (Thermo Fisher Scientific), coupled online with an Orbitrap Fusion Lumos (Thermo Fisher Scientific). Chromatographic separation was performed with a two-column system, consisting of a C-18 trap cartridge (300 µm ID, 5 mm length) and a picofrit analytical column (75 µm ID, 25 cm length) packed in-house with reversed-phase Repro-Sil Pur C18-AQ 3 µm resin. Peptides were separated using a 30 min gradient from 1-30% buffer B (buffer A: 0.1% formic acid, buffer B: 80% acetonitrile + 0.1% formic acid) at a flow rate of 300 nl/min. The mass spectrometer was set to acquire spectra in a data-independent acquisition (DIA) mode. Briefly, the full MS scan was set to 300-1100 m/z in the orbitrap with a resolution of 120,000 (at 200 m/z) and an AGC target of 5x10^5^. MS/MS was performed in the orbitrap with sequential isolation windows of 50 m/z with an AGC target of 2x10^5^ and an HCD collision energy of 30.

Histone peptides raw files were imported into EpiProfile 2.0 software^65^. From the extracted ion chromatogram, the area under the curve was obtained and used to estimate the abundance of each peptide. To achieve the relative abundance of post-translational modifications (PTMs), the sum of all different modified forms of a histone peptide was considered as 100% and the area of the specific peptide was divided by the total area for that histone peptide in all of its modified forms. The relative ratio of two isobaric forms was estimated by averaging the ratio for each fragment ion with different mass between the two species.

### Negative contrast staining and immuno-transmission electron microscopy

EVs were visualized by negative staining as described previously by Hedlund *et al.*^66^. A total of 4 µL of sample was adsorbed to glow discharged, formvar/carbon coated copper grids. Following two washes in ddH_2_O, negative contrast staining was performed in 1.5% (w/v) uranyl acetate for 30 s and dried at RT. Immuno-TEM on EVs was performed as previously described^67^ with slight modification. Samples were absorbed on to copper grids and blocked by 0.1% (w/v) gelatin and 1% (w/v) BSA in PBS for 1 h at RT. After brief washing in PBS, grids were incubated in primary antibodies (see the Antibodies section, above) diluted in PBS containing 1% BSA and incubated overnight at 4°C. Grids were washed in PBS and incubated with 10 nm protein A-gold particles (PAG10, BBI Solutions) for 1 h at RT. After washing using PBS, grids were washed 5 times in water and exposed to 1.5% uranyl acetate for 30 s and dried at RT before imaging.

Immuno-TEM on cells was performed according to published methods^68^. OLN-93 cell sections were processed according to the Tokuyashu method^69^. Cells were fixed in freshly prepared EM fix solution containing 2% (w/v) paraformaldehyde and 0.2% (w/v) glutaraldehyde. Optimal fixation was facilitated using a microwave incubator (PELCO BioWave). Cells were then centrifuged for 5 min at 500 xg and the fixative solution was replaced with PBS containing 10 mM glycine to quench free aldehyde groups. Thereafter, 3x PBS washes were performed and cells were resuspended in 12% (w/v) gelatin and incubated for 1 h at 37°C. Gelatin blocks were placed into 2.3 M sucrose overnight at 4°C on a rocking platform. Blocks were mounted on the top of metal pins and snap frozen in liquid N_2_. Cryo-sectioning (70 nm sections) was performed in a Leica UC7 cryostat (Leica Microsystems), and sections were mounted on copper 300x300 grids. Grids bearing several sections were placed in PBS and incubated at 37°C for 1 h followed by incubation in 10 mM glycine in PBS. After 3x washes in PBS the sections were blocked in 1% (w/v) BSA, 0.1% (w/v) gelatin. Primary antibodies for histone H1 (most sensitive for this method) and CD63 were added and incubated overnight at 4°C. After 5x washes in PBS, secondary antibody-gold or protein-A gold conjugates were added. Finally, sections were washed extensively in PBS and contrast enhanced with uranyl acetate/methylcellulose (1:4) solution. Control sections processed without primary antibodies were prepared in parallel to estimate the specificity and background of antibody binding. Specimens were examined with a JEM1230 (Jeol) or Talos electron microscope (Thermo Fisher Scientific) at 80 kV and micrographs were collected with a MSC 600CW CCD camera (Gatan) using DigitalMicrograph software (Gatan).

### Statistics

Statistical analyses were performed using Graph Pad Prism 5.0c (GraphPad Software) or SPSS 26 (IBM). Error bars represent the standard error of mean. Data was assessed for normality using Shapiro-Wilk test. Results were analysed for statistical significance by T-test, one-way or two-way analysis of variance (ANOVA) followed by the Bonferroni and Sidak’s post hoc test. Treatment effects were assessed using paired sample T-test.

## RESULTS

### Histone association with extracellular and intraluminal vesicles

First, we characterized histone secretion into the conditioned media from the rat oligodendrocyte progenitor cell line OLN-93. This line could be grown in 2D culture under serum-free conditions for several days, facilitating purification of EVs by sequential centrifugation ^47^. In addition, it was possible to produced large numbers of EVs in 3D culture in a bioreactor, enabling us to generate sufficient material for large scale biochemical characterization.

Comparison of histones in the whole cell lysate (WCL) and EV pellet fraction (EV^100,000^ ^xg^) by western blotting (WB) showed significant enrichment of membrane localized tetraspanins CD9 and CD63 and luminal markers tumor susceptibility gene 101 (Tsg101) and actin beta (Actb) in EVs (Fig. 1A). Lack of contamination of with cellular organelles was confirmed by absence of the cis-Golgi matrix protein GM130. As expected, robust signals were observed for all nuclear core histones (H2A, H2B, H3 and H4) in the WCL, but also in EVs. We then used immunogold staining together with transmission electron microscopy (immuno-TEM) to confirm that all histones (core and linker) localized to EVs. Surface labelling in unpermeabilized EVs suggested that like CD63 (Fig. 1B) and CD9 (Fig. S1A), histones also localized to the EV surface (Fig. 1B and S1A)

**Figure 1.**
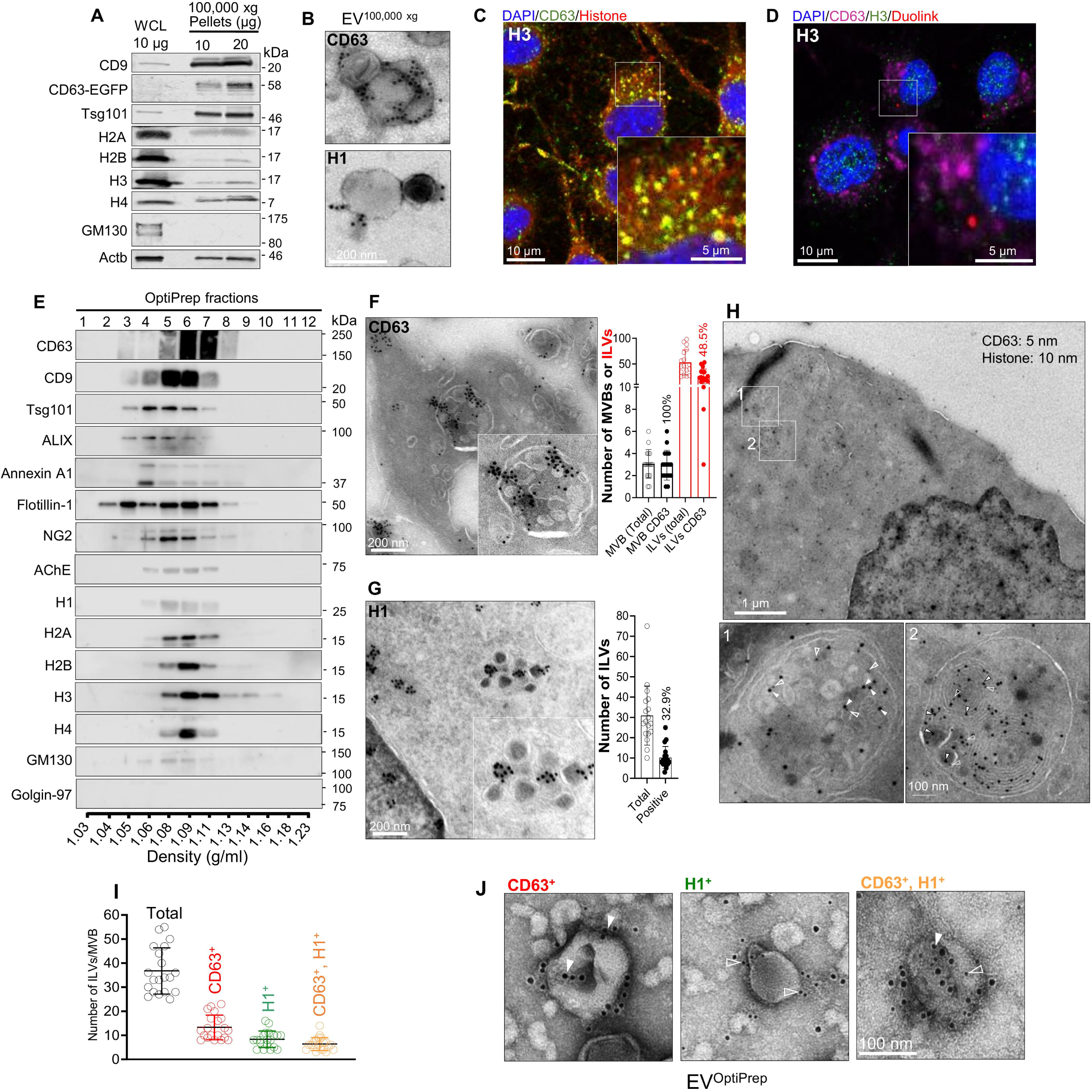
Histones are associated with ILVs and EVs. **(A)** WB of whole cell lysate (WCL, 10 μg) from a stable OLN-93 cell line expressing human CD63-EGFP and EVs (10, 20 μg) from 100,000 xg pellet. CD9, CD63, Tsg101, core histones (H2A, H2B, H3 and H4), Golgi marker GM130 as well as Actb identified using specific antibodies. Approximate molecular mass is shown in kDa. **(B)** Immuno-TEM micrographs of OLN-93 EV^100,000^ ^xg^ using anti-CD63 primary antibody and secondary antibody conjugated to with 10 nm gold. **(C)** Confocal micrographs from a single z-plane of OLN-93 cells showing colocalization of anti-H3 (red) with anti-CD63 (green). Nuclei were counterstained with DAPI (blue). Inset shows a cropped and expanded region of the main image. **(D)** Confocal micrograph showing Duolink staining (red) due to colocalization of anti-H3 (rabbit) and anti-CD63 (mouse) antibodies. **(E)** WB showing localization of a range of membrane, EV and organelle markers together with linker and core histones in OptiPrep purified EV^100,000^ ^xg^ (300 µg) fractions from a 12%-36% (w/v) equilibrium density gradient. One 20^th^ of each fraction was loaded on the gel. Experiments were repeated 3 times, and one representative image is shown. **(F)** Representative immuno-TEM micrograph of OLN-93 cell sections showing anti-CD63. Inset panel shows a magnified region containing an MVB. The column chart on the right shows quantification of the total number of MVBs versus CD63^+^ MVBs (black), and total number of ILVs versus CD63^+^ ILVs (red) as % of total. **(G)** as for (F) but showing anti-H1. Column chart on the right shows the total number and H1-positive ILVs per MVB. Error bars represent the mean ±SD. **(H)** Immuno-TEM showing differential labelling of anti-CD63 (5 nm gold, open arrowheads) and anti-H1 (10 nm gold, solid arrowheads). Panels below (1 and 2) show higher magnification images of the regions indicated in H. Solid arrowheads indicate anti-H1 labelling and open arrowheads indicate anti-CD63 labelling. **(I)** Scatter plot showing quantification of CD63 and H1 labelling and colocalization (on the same ILV). A total of 10 images from separate cells and sections were quantified and the numbers of ILVs per MVB are represented by each point. Error bars represent the mean ±SD. **(J)** Immuno-TEM micrographs of OLN-93 EV^OptiPrep^ showing anti-CD63 (10 nm gold, solid arrowheads) and anti-H1 (5 nm gold, open arrowheads). Note gold particle sizes are opposite to those used in (F).

To see whether histones colocalized with CD63 in intact cells, we first used immunofluorescence staining in combination with confocal microscopy. Colocalization of non-nuclear staining for all four core histones and CD63 was observed in cytoplasmic or membrane puncta (Fig. 1C, Fig. S1B-D). Since the degree of colocalization observed was greatest with H3/CD63 (Fig. 1C), we also tested this antibody combination in a proximity ligation assay (Duolink PLA), which has a higher spatial resolution (within 40 nm) than standard confocal fluorescence microscopy. PLA confirmed an association of anti-H3/CD63 antibodies in cytoplasmic puncta with the morphological appearance of MVBs (Fig. 1D). No background signal was observed in the absence of primary antibodies (Fig. S1E) confirming the specificity of the assay.

OptiPrep discontinuous density gradient centrifugation is a well-established method for the purification of small EVs (sEVs) including smaller MVs and exosomes, from other components such as larger microvesicles and apoptotic bodies^70, 71^. In EVs from mast cell and erythroleukemic cell lines, histones localize to high denisty (HD; 1.14-1.19 g/ml) but not to low density (LD; 1.10-1.13 g/ml) sEV fractions^30^. To investigate histone distribution in EVs from OLN-93 cells, we fractionated EV^100,000^ ^xg^ on 12-36% OptiPrep gradients. Fraction densities were calculated based on a reference gradient (Fig. S2A). TEM was used to confirm the purity and morphology of EVs in each fraction (Fig. S2B). The majority of EVs with an intact morphology and ranging from 30-150 nm localized to fractions 3-7 (Fig. S2B). WB confirmed enrichment of CD63 in fractions 3-8 (1.05-1.13 g/ml) and most abundantly in fractions 6-7 (1.09-1.11 g/ml, Fig. 1E). CD9, Tsg101, and ALG-2-interacting protein X (ALIX/PDCD6IP) showed shifted distributions in comparison to CD63, being enriched in fractions 3-6 (Fig. 1E). Annexin A1, a marker of plasma membrane-derived microvesicles, was most abundant in fraction 4 (1.06 g/ml). Flotillin-1 (membrane marker) spanned fractions 2-7, whereas extracellular acetylcholinesterase (AChE) and the NG2 chondroitin sulfate proteoglycan, which is a membrane-anchored marker of oligodendrocyte progenitors, localized to fractions 4-7. The Golgi marker GM130 was seen at low levels through fractions 3-9, but Golgin-97 was not detected (Fig.1E). Thus, we were able to resolve distinct but overlapping EV populations ranging from Annexin A1 positive microvesicles (fraction 4), through Tsg101/Alix positive sEVs in fractions 4-5 and CD9/CD63 tetraspanin positive sEVs in fractions 5-7. Interestingly, in OLN-93 cells all four core histones and H1 localized to fractions 4-7 and were most abundant in fraction 6, identifying them as components of sEVs that co-purify with CD63, CD9, Tsg101, and Alix (Fig.1E).

To investigate the distribution of histones in EVs from other cell lines, we fractionated EV^100,000^ ^xg^ pellets from a selection of mouse and human cells with different tissue origins using OptiPrep gradients. H3 positive LD and HD fractions were identified for all cell lines tested (HeLa, Caco-2, NSC-34, AC-16 and A549; Fig. S3-S4) though with overlapping distributions. H3 was found in fraction 8 (1.13 g/ml) in all lines tested and TEM analysis confirmed the presence of EVs in histone positive fractions. Interestingly, HeLa EVs resolved as two distinct H3 containing populations (Fig. S3; fraction 3-4 copurifying with Alix and Tsg101 and fractions 6-8). Hence, H3 seems to localize to EVs that float over 3 different density ranges – LD fractions 3-4 (mainly Tsg101+, Alix+; from HeLa and to a lesser extent from Caco-2), HD fractions 5-8 (tetraspanin positive; all cell lines apart from A549) and fractions 9+ (OLN-93, NSC-34, AC-16, A549).

EV^100,000^ ^xg^ and EV^OptiPrep^ fractions may contain a mix of plasma membrane derived microvesicles (indicated by the aforementioned presence of Annexin A1 in fractions 4-8; Fig. 1E) and exosomes that form as intraluminal vesicles (ILVs) in MVBs. To see if histones could be identified in association with ILVs, we developed a protocol for immuno-TEM on ultrathin sections of OLN-93 cells. All MVBs examined contained CD63 positive ILVs (49%, ranging from 3-50 per MVB, Fig. 1F). The densest histone labelling was observed for the histone H1 antibody where positive ILVs were identified in 32.9% of cases (Fig. 1G). We also confirmed labelling for core histones (24-41% of ILVs; Fig. S5A-D).

We used immuno-TEM with secondary antibodies conjugated to differently sized gold particles to verify the colocalization of histone and CD63 in MVBs. OLN-93 sections were probed with anti-H1 (10 nm gold) and anti-CD63 (5 nm gold; Fig. 1H) in combination. Quantification showed 36% of ILVs labelled positively for CD63 and 23% for H1. Interestingly, 17% of ILVs co-labelled with both anti-CD63 and anti-H1 (Fig. 1I). We also confirmed the colocalization of CD63 and H1 on EV^OptiPrep^ fractions 4-7 (Fig. 1J). The presence of histone in ILVs and EV^OptiPrep^ from human HeLa (Fig. S6A-B) and Caco-2 cell lines (Fig. S7A-B) was also confirmed. In summary, using a range of methodological approaches, we have shown that histones are associated with ILVs inside MVBs and on secreted EVs in several different cell lines. A proportion of these ILVs/EVs are positive for both histones and CD63.

### Histone secretion in response to cell stress

Cellular stress is a common denominator in pathological conditions where elevated circulating histones have been observed^72^ and we hypothesized that stress might enhance histone secretion via EVs. Therefore, we developed a heat stress paradigm whereby we harvested/changed the conditioned media on OLN-93 cells as a control condition (‘Before’) and then cultured the cells at 42°C for 3 h, followed by a 4 h recovery period at 37°C (‘Heat stress’). Media was collected, changed, and harvested at a third time point after 17 h (‘After’; Fig. 2A).

**Figure 2.**
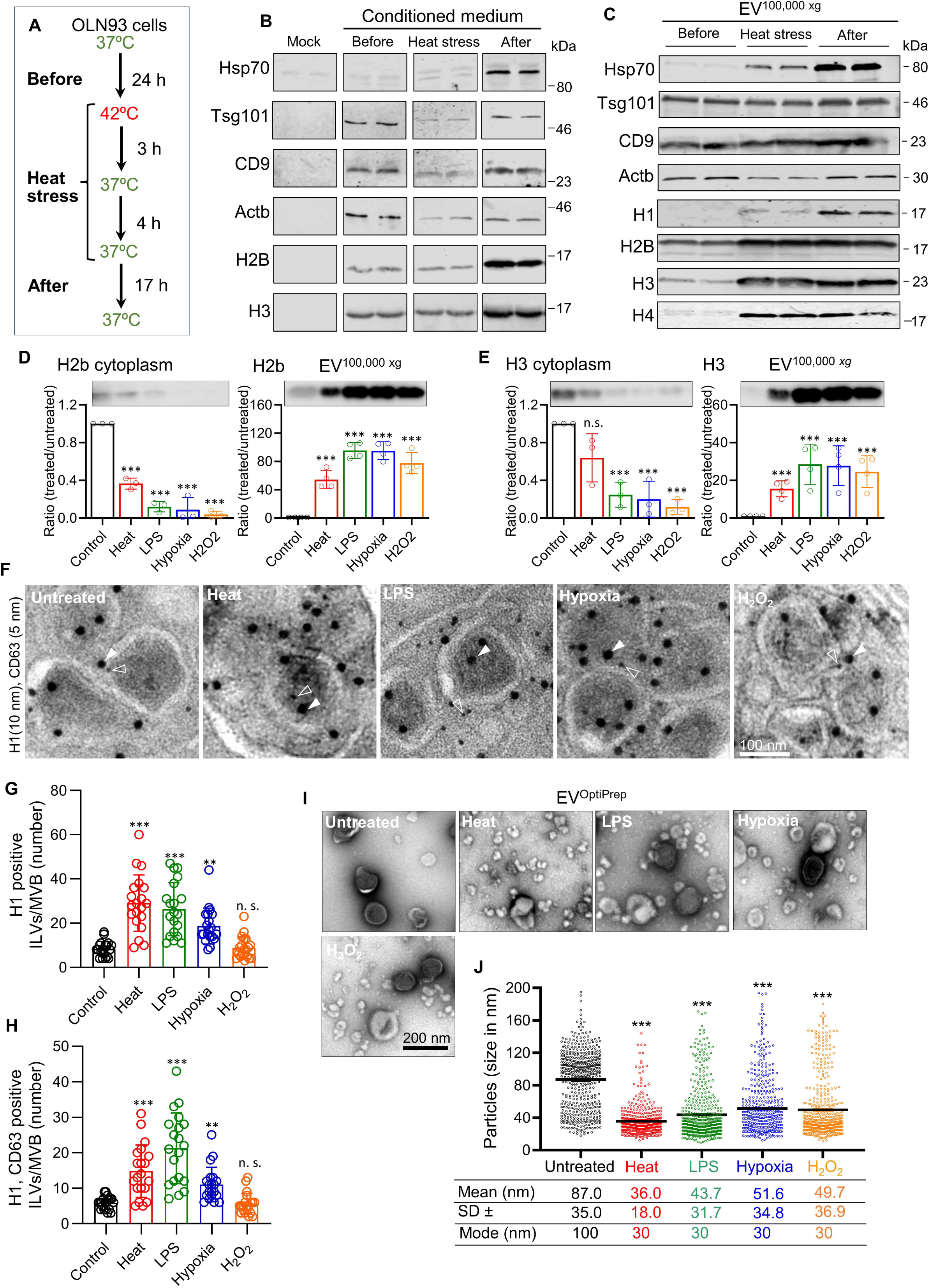
Cellular stress enhances the release of histones via ILVs/EVs. **(A)** Schematic diagram outlining the experimental procedure for the heat stress experiment, showing sample collection times for Before, Heat stress and After conditions. **(B)** WB showing markers (specific antibodies as shown) detected after pulldown with StrataClean resin of total protein from 1 ml of conditioned media from Before, Heat stress and After samples, as well as fresh, unconditioned media (Mock). **(C)** WB of EV^100,000^ ^xg^ (11 µg per lane) isolated from the same conditioned media samples shown in (B). **(D-E)** OLN-93 cells were stressed by heat, LPS, hypoxia, and H_2_O_2_ followed by WB to quantify histone H2B and H3 in cytoplasm and EV^100,000^ ^xg^. Column charts show the ratio between band intensities for stress versus control conditions. Three independent experiments were performed, and one representative WB image is shown. **(F)** Immuno-TEM images of control and stressed OLN-93 cell sections showing ILVs within an MVB. Anti-CD63 (5 nm gold; open arrowheads) and anti-H1 (10 nm gold; closed arrowheads). **(G)** Quantification of number of H1 positive ILVs per MVB present under the various conditions. **(H)** Co-localisation of anti-CD63/H1 on the same ILV under control and stress conditions. Error bars represent the SD. One-way ANOVA was used to compare control and stress conditions. *, *p* ≤ 0.05, ***, *p* ≤ 0.001. **(I)** Representative TEM micrographs of negatively stained OLN-93 EV^OptiPrep^ isolated from control and stress conditions as indicated in each panel. **(J)** Scatter plots showing size distribution of EVs (n = 500 EVs measured from n=10 randomly selected images) measured in TEM micrographs from each condition. Statistical significance between control and stress conditions (***, *p* ≤ 0.001) using one-way ANOVA followed by Bonferroni’s multiple comparison post-test. Horizontal bars in each plot represent the mean diameter in nm, shown below with the ±SD, as well as the mode size (in 10 nm bins).

To quantify secreted histones, we first pulled down all proteins from 1 ml of conditioned media using StrataClean resin and analysed by WB. We observed strong induction of the heat shock protein Hsp70 in the After sample, confirming the efficacy of the treatment (Fig. 2B). EV markers Tsg101, CD9 and Actb showed a slight reduction in the Heat stress sample, likely due to the shorter period of media conditioning (7h versus 24 h for Before). However, levels of H2B and H3 were equivalent in Before and Heat stress samples and increased in the After sample (Fig. 2B), suggesting elevated secretion of histones in response to heat stress. These findings were confirmed in EV^100,000^ ^xg^ fractions where a constant amount of protein was analysed by WB to account for the different periods of media conditioning. Here, we observed strong induction of Hsp70 in both Heat stress and After samples (Fig. 2C). The levels of Tsg101, CD9, and Actb remained constant. However, heat stress resulted in increased secretion of linker and core histones. Hence, EV associated histones accumulate in the conditioned media and are increased in EVs in response to heat stress.

To see whether histone secretion is part of a common stress response, we exposed OLN-93 cells to several stress paradigms. EVs were harvested after 24 hours, though the period of exposure to the stressor was different in each case (see Methods): Heat (42°C for 3 h), the endotoxin (LPS; 24 h), low oxygen tension (hypoxia; 24 h), and oxidative stress (H_2_O_2_; 24 h). Since histones released from dead/dying cells could complicate analyses, we first determined the effect of the stresses on cell viability using live/dead (PI/Hoechst 33342) staining, and on apoptosis using Annexin V/Cytox staining. A significant increase in cell death, indicated by nuclear PI labelling, was observed after heat stress (Fig. S8A, B) but not after the other stress paradigms. Similarly, a significant increase in Annexin V positive (early apoptotic), Cytox positive (compromised plasma membrane, as for PI) and Annexin/Cytox double positive (late apoptotic/dead) cells were observed following heat stress, but not for other stress conditions (Fig, S8C, D). Interestingly, LPS treatment led to reduced Annexin V, Cytox and double stained cells in comparisons to controls, indicating a potential protective effect of LPS exposure in these cultures (Fig. S8D).

We then proceeded to quantify the relative levels of histones in cytoplasmic versus EV^100,000^ ^xg^ fractions under control and stress conditions. All stress interventions led to a decrease in the cytoplasmic pool of H2B and H3 of between 10-64% (Fig. 2D, E). Loading controls for these blots are shown in figure S9. In contrast, large increases in H2B and H3 were found in the EV^100,000^ ^xg^ fractions under all stress conditions ranging from 15-95-fold (Fig. 2D, E). To validate the results of WB in intact cells, we used immunostaining followed by confocal microscopy. Enhanced colocalization of anti-H3 and anti-CD63 staining was observed in cytoplasmic MVB-like compartments under all stress conditions (Fig. S10A). In contrast, colocalization with the plasma membrane/microvesicle marker Annexin A1 appeared unchanged (Fig. S10B). Duolink staining with anti-H3 and anti-CD63 showed enhanced colocalization of these 2 markers under all stress conditions (Fig. S10C). Further, immuno-TEM was used to quantify colocalization of histone and CD63 more precisely. Increased numbers of anti-H1 labelled (10 nm gold) and anti-CD63 labelled (5 nm gold) particles were found in the MVBs of cells exposed to all stress conditions (Fig. 2F, S11A-E). The number of H1 or H1/CD63 double positive ILVs increased significantly under all stress conditions apart from oxidative stress (Fig. 2G-H), which may reflect the short half-life of the stressor (see Methods) and/or the longer recovery period available to cells subjected to this stress paradigm.

To investigate if the size and quantity of EVs changed in response to stress, first we analysed EV^100,000^ ^xg^ from Before, Heat stress and After by NTA (Fig. S12A). There was an approximately 3-fold increase in EV particle concentration in Heat stress and After samples after normalization for the conditioning period (Before = 2.6x10^7^, Heat stress = 7.1x10^7^, After = 8.6x10^7^ particle/ml/h; Fig. S12B). We also observed an increase in 40-100 nm sized particles (Fig. S12A). However, since the sensitivity of NTA declines for particles of less than 50 nm^30^, we used TEM to estimate the size of EV^OptiPrep^. All stress conditions led to the secretion of smaller diameter EVs (mean = 36-52 ±18-37 nm, mode = 30 nm; Fig. 2I-J) in comparison to untreated samples (mean = 87 ±35 nm, mode = 100 nm; Fig. 2J). In summary, our results show that cellular stress responses lead to an increase in the quantity and a reduction in the size of EVs.

### Co-dependence of histones and DNA for EV binding

Previous studies have focussed on the DNA/chromatin content of sEVs and concluded that sEV histones are tightly associated with DNA as nucleosomes. However, the situation is unclear since both double- and single-stranded (ds- or ss-, respectively) DNA have been found inside and outside of EVs^40, 73^. As only dsDNA associates with histones in nucleosomes, we sought to clarify both the possible co-dependence of histones and DNA for EV binding, and their location in relation to EV membrane and lumen.

First, we quantified the amount of DNA (Qubit) and H3 (WB) in EV^OptiPrep^ fractions from OLN-93 cells cultured under control or stress conditions. H3 showed a dynamic distribution, depending on the type of stress and period of exposure to the stressor (Fig. 3A). Loading controls of the blots are shown in figure S13. H3 increased in fractions 6-7 under all stress conditions but DNA did not, with the majority of DNA/µg total protein localizing to fractions 8-12. Notably, the distributions of DNA did not mirror the distributions of H3 suggesting a limited co-dependency of histones and DNA for EV binding.

**Figure 3.**
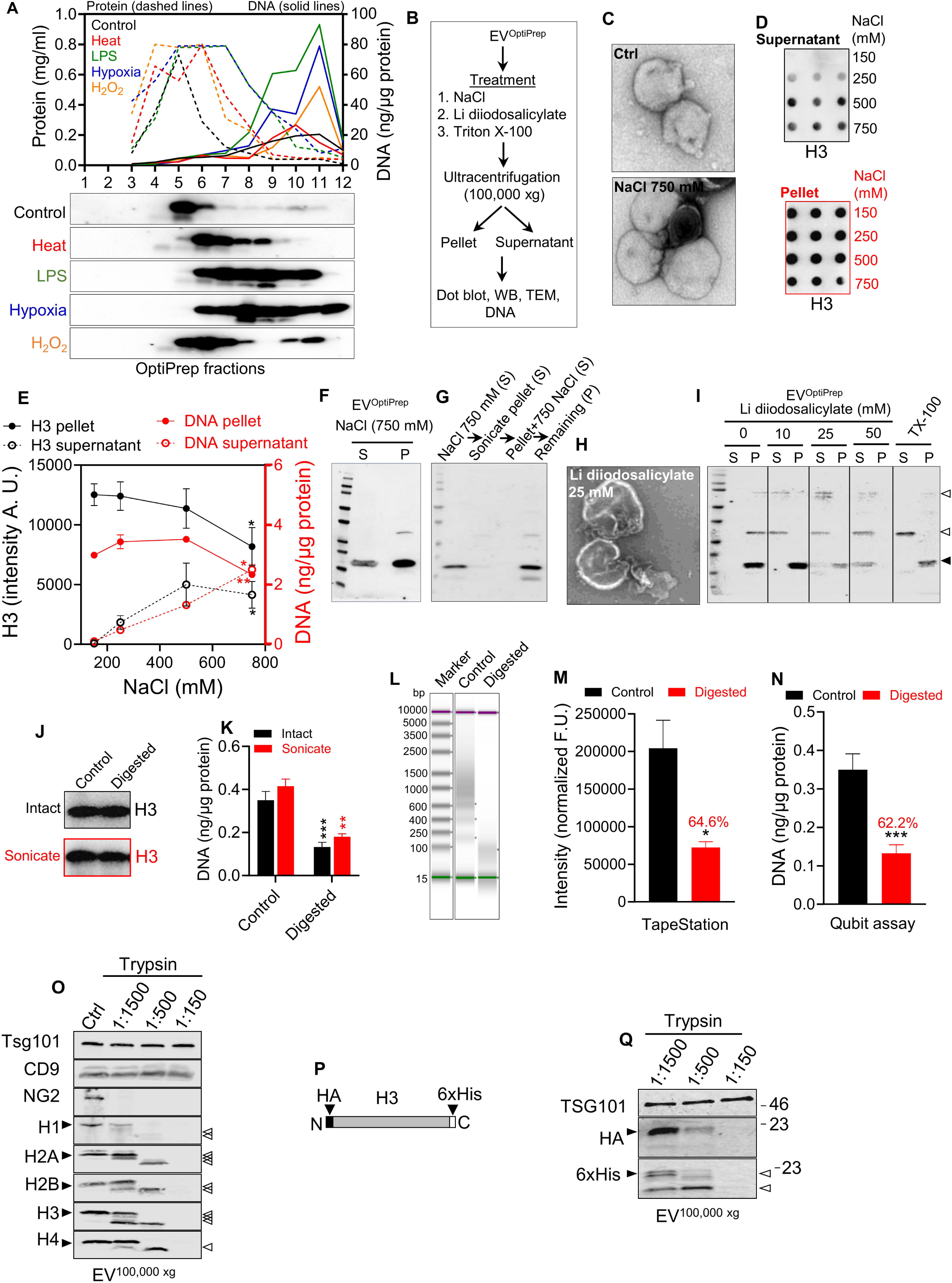
Histones associate with the EV membrane. **(A)** Line graphs (above) showing quantification of protein (dashed lines) and DNA (solid lines) by Qubit in OptiPrep fractions from different stress conditions. WBs (below) showing distribution of H3 in OptiPrep fractions. **(B)** Schematic diagram showing the basic procedure used to extract histones from EV^OptiPrep^. **(C)** TEM images showing control (Ctrl) EVs or following 750 mM NaCl treatment. **(D)** Extraction of histones from OptiPrep purified (fractions 4-7; EV^OptiPrep^). Approximately 10 µg EVs were treated with 150-750 mM NaCl for 24h at 4°C, followed by ultracentrifugation and estimation of H3 in the supernatants and pellets by dot blot. **(E)** Line graphs showing the intensities of dot blots (H3 quantification, black) and DNA content (Qubit assay). Statistical analysis between 150 mM NaCl as control and other NaCl concentrations by one-way ANOVA, *, *p* ≤ 0.05, **, *p* ≤ 0.01. **(F)** Extraction of histones in the supernatant (S) and remaining in the pellet (P) from EV^OptiPrep^ treated with 750 mM NaCl and shown by WB. **(G)** S and P fractions of EV^OptiPrep^ after extraction with 750 mM NaCl, then sonication and re-pelleting, followed by a further round of extraction with 750 mM NaCl to remove luminal contents. The remaining pellet shows residual H3 that was not extracted by 750 mM NaCl. **(H)** TEM images showing intact EV membranes after 25 mM lithium 3,5-diiodosalicylate treatment. **(I)** Extraction of histones into the supernatant (S) or remaining in the pellet of EV^OptiPrep^ following treatment with increasing concentrations of lithium 3,5-diiodosalicylate (10-50 mM) or Triton X-100 (0.1% TX-100). **(J)** WB showing H3 present in EV^OptiPrep^ (control or sonicated) with and without nuclease digestion (DNAseI, S1 nuclease and Exosnuclease III). **(K)** Column chart showing DNA quantification (Qubit) in same samples as (J). **(L)** TapeStation gel image of total EV DNA purified from control and digested samples. **(M)** Column chart showing quantification of DNA by TapeStation and, **(N)** Quantification of DNA in same samples using Qubit. **(O)** Analysis of OLN-93 EV^OptiPrep^ by WB without (Ctrl) or following trypsin treatment (1:1500-1:150 μg trypsin:μg EV protein). Presence of truncated bands for histone proteins following trypsin treatment are indicated on right by open arrowheads. **(P)** Schematic diagram of epitope tagged H3 cassette used to express the protein in a HEK293 stable cell line. HA-tag is located at the N-terminus and a 6xHis-tag at the C-terminus (not to scale). **(Q)** WB analysis of EV^100,000^ ^xg^ isolated from stable HA-H3-6xHis HEK293 cell line after trypsin treatment. Location of full-length HA-H3-6xHis, as well as partially degraded species are indicated by a black or open arrowheads, respectively. Control recombinant histone digestion by trypsin is shown in supplementary Fig. S12.

Next, we devised a protocol combining chemical extraction, ultracentrifugation, and sonication to distinguish between histones or DNA loosely associated with the outer surface of EVs from those tightly bound to, or integrated into, the membrane, or in the lumen (Fig. 3B). A high concentration of NaCl is commonly used to disrupt electrostatic interactions whilst still being a non-denaturing salt^58^. The chaotropic salt lithium 3,5-diiodosalicylate can solubilize membrane associated proteins at lower concentrations, while disrupting the membrane completely at higher concentrations^59^. In addition, we used the detergent TX-100 (0.1%) to lyse EVs^74^, or sonication to disrupt them. TEM was used to estimate EV morphology and integrity. Following EV re-isolation via ultracentrifugation, we quantified H3 and DNA in the supernatant or pellet fractions using WB or Qubit/Tape Station, respectively.

EV^OptiPrep^ could be treated with up to 750 mM NaCl without adverse effect on morphology (Fig. 3C, S13A). However, 1M NaCl led to visible damage to EV membranes (Fig. S14A). Increasing concentrations of NaCl (250-750 mM) led to a displacement of approximately 40% of H3 into the supernatant, which became maximal at 500 mM NaCl (Fig. 3D-E, and S13). Up to 30% of EV associated DNA was displaced but did not reach a plateau, even at 750 mM NaCl (Fig. 3E).

A significant proportion of H3 remained associated with the EV pellets following treatment with 750 mM NaCl (Fig. 3F). To probe this in more detail, EV pellets recovered after 750 mM NaCl treatment, were sonicated to disrupt membranes, and expose luminal contents. We observed that a minor proportion of H3 was released by this procedure, but the majority remained associated with the pellet and could not be extracted with an additional 750 mM NaCl treatment of sonicated EVs (Fig. 3G). Interestingly, following different stress conditions a variable proportion of surface associated H3 (extracted by 750 mM NaCl) could be removed. Heat stress EVs showed the least surface associated H3, while LPS and hypoxia EVs showed the highest levels (Fig. S14B).

We used lithium 3,5-diiodosalicylate extraction to estimate the proportion of H3 that was tightly associated or integral to the membrane. TEM analysis showed that EV membranes remained largely intact in 25 mM lithium 3,5-diiodosalicylate (Fig. 3H) but were disrupted by 50 mM (Fig. S14C). H3 was extracted progressively into the supernatant by 10-25 mM and completely by 50 mM lithium 3,5-diiodosalicylate. TX-100 treatment led to extraction of approximately 50% H3 into the supernatant (Fig. 3I). Taken together, our results suggests that a significant proportion (50-60%) of EV histones are tightly associated with the EV membrane. Some histones associate more loosely with the surface of EVs, likely via electrostatic interaction, and a minor fraction may be luminal.

We treated EV^OptiPrep^ with a combination of nucleases (DNase I, S1 nuclease and Exonuclease III) to remove DNA from EVs. In addition, we sonicated EVs to expose luminal contents. Digestion had no appreciable effect on H3 association in both intact and sonicated EVs (Fig. 3J). However, based on Qubit estimation in the same samples, the amount of DNA was significantly reduced and sonication had no additional effect (Fig. 3K). To determine DNA size and amount more precisely, we purified EV DNA and analysed this by TapeStation and Qubit in parallel. Nuclease digestion led to the degradation of EV DNA to ≤100 bp (Fig. 3L), with ≈ 40% remaining (Fig. 3M), which was confirmed by Qubit (Fig. 3N).

We used a range of trypsin concentrations to probe exposure on the surface of EVs and analysed histones and other proteins in the pellet by WB (Fig. 3O). Under conditions where the membrane associated proteins Tsg101 and CD9 were protected from digestion, a complete loss of the glycophosphatidylinositol (GPI)-linked outer membrane NG2 occurred even at the lowest trypsin:protein ratio (1:1500). All histones were partially protected from trypsin cleavage at 1:1500-1:500, although they were completely removed at 1:150. Recombinant purified histones were used as control for trypsin digestion and, in contrast to EV associated histones, did not show a similar pattern of partial cleavage products (Fig. S15).

Since all anti-histone antibodies tested were directed against C-terminal epitopes (see Methods) we also examined whether N-terminal of C-terminal regions were equally susceptible to cleavage in intact EVs. A HEK293 cell line stably expressing H3.1 with an N-terminal HA- and a C-terminal 6xHis-tag (Fig. 3P) were generated. EVs were purified and treated with trypsin as above. With an increasing trypsin:protein ratio, reactivity towards the N-terminal HA-tag was lost without the appearance of smaller proteolytic fragments whereas the C-terminal 6xHis antibody recognized one or more truncated species (Fig. 3Q). In summary, our results show that histones are mainly EV membrane associated proteins. Furthermore, although DNA is also present at the surface of EVs, the majority of histone does not appear to be dependent on DNA for its membrane association.

### Binding of histone octamers and nucleosomes to membrane lipids and EVs

To probe the potential interaction of histones with lipids, we prepared liposomes from OLN-93 whole-cell lipids. After incubation with recombinant human histones or purified total histones from calf thymus, liposome-histone mixtures were washed using 100,000 kDa molecular weight cut-off ultrafiltration and analysed by WB. All histones bound to liposomes and were retained by the filter even after thorough washing (Fig. 4A). Liposome-histone mixtures were also resolved on 0-40% sucrose gradients^75^. Total histones from calf thymus, or individual recombinant histones floated to lower density fractions and were found in the least dense fractions 1-2 (Fig. 4B). These experiments confirm that recombinant histones expressed in *E.coli*, which lack PTMs, bind directly to membrane lipids.

**Figure 4.**
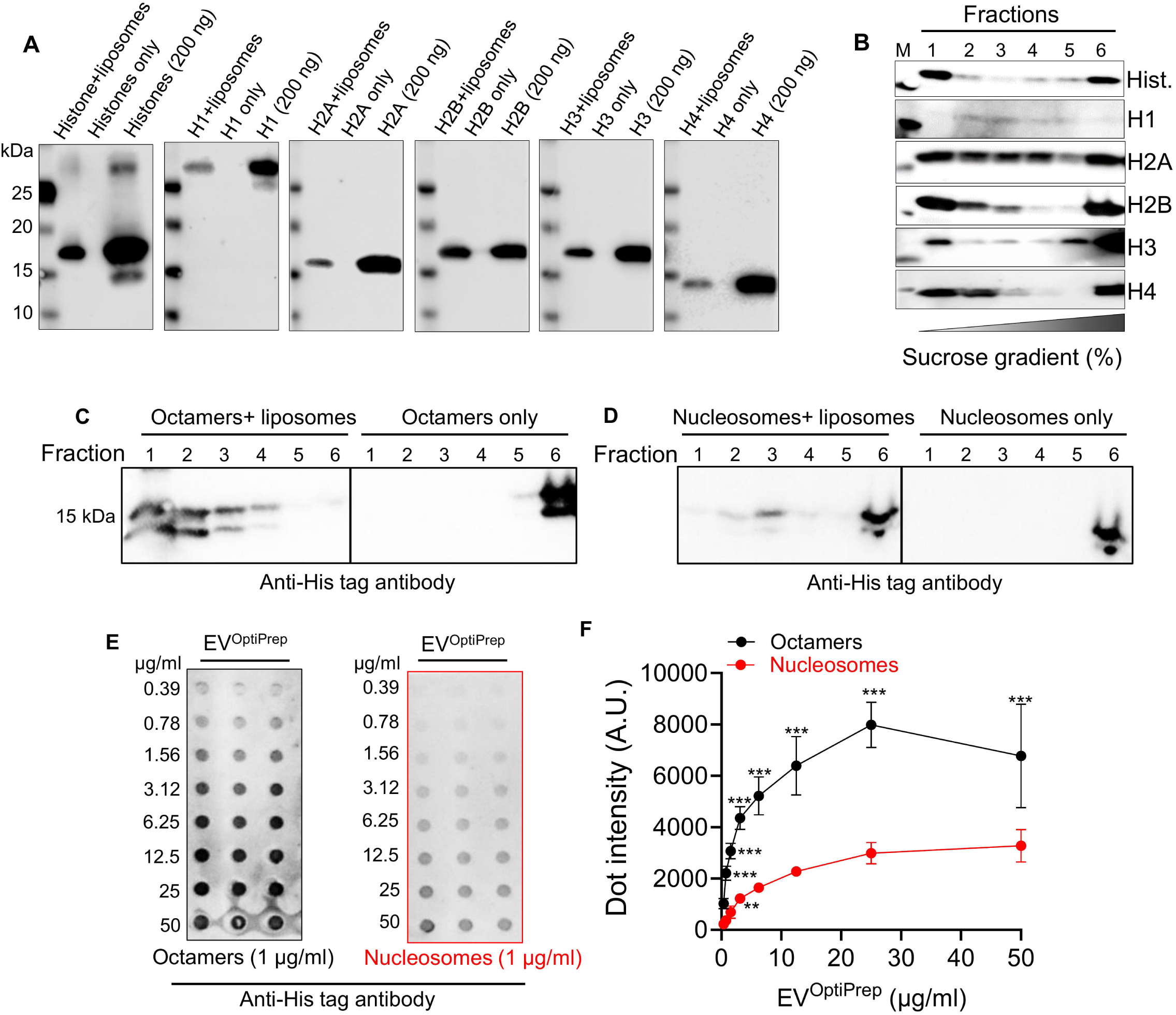
Preferential membrane binding of histones and octamers versus nucleosomes. **(A)** Analysis of histone interaction with liposomes. Total histones or individual recombinant histones (as indicated) were mixed with liposomes derived from OLN-93 total cellular lipids and washed extensively in a 100,000 MWCO spin concentrator. The retentate was subjected to WB with specific anti-histone antibodies. A histone protein-only control (without liposomes) was used to control for lipid-independent retention of histones by the MWCO filter. Histone proteins (200 ng as indicated) were loaded as references. **(B)** Liposome flotation on a sucrose density gradient (0-40%). Histones as in (A) were mixed with liposomes and then added to the bottom layer (40% sucrose) of the gradient and subjected to ultracentrifugation at 100,000 xg. Fractions were collected and analysed by WB. **(C)** Recombinant histone octamers (5 µg) proteins were mixed with liposomes (20 µg) and separated as in (B). Octamers were detected by WB using anti-His tag antibody. **(D)** As for (C) but using recombinant nucleosomes instead of octamers. **(E)** Dot blot showing binding of octamers or nucleosomes to EV^OptiPrep^. Serially diluted EVs were blotted onto a nitrocellulose membrane and probed with 1 µg/ml octamers/nucleosomes. The bound fractions were detected using an anti-His tag antibody. **(F)** Line graphs showing the intensity of dot blots plotted from (E). Two-way ANOVA and Bonferroni multiple comparison test was performed: **, *p* ≤ 0.01, ***, *p* ≤ 0.001,

Next, to test if DNA enhances the binding of histones to liposomes, we used sucrose density gradient floatation to compare the relative binding of recombinant histone octamers and mono nucleosomes. His-tagged octamers/nucleosomes alone remained in fraction 6 (Fig. 4C, D), whilst most of the added histone octamer bound to liposomes and floated in fractions 1-2 (Fig. 4C). In contrast, nucleosomes bound poorly to liposomes (Fig. 4D). EVs did not float reliably on sucrose gradients when treated with nucleosomes (not shown), so we investigated EV binding using a solid phase (dot blot) assay. First, we immobilized different amounts of EVs (0.39-50 μg/ml) on a nitrocellulose membrane, then incubated with His-tagged octamers/nucleosomes, followed by detection of binding with an anti-His tag antibody. Results showed that octamers bound significantly higher to EVs in comparison to nucleosomes, indicating the presence of DNA in the nucleosome interferes with histone/octamer interaction to EV membranes. (Fig. 4E, F, and S13).

### Origin of extracellular vesicle associated histones

Histones could localize to EVs via several different pathways: i) Sorting of soluble cytoplasmic histones into MVBs via ESCRT-dependent or -independent pathways, thereby partitioning to the luminal EV compartment. ii) Acquisition of histones to the outer surface of EVs via the amphisome pathway, following fusion of autophagosomes to endosomes. iii) Endocytosis of EV or non-EV derived histones and their subsequent sorting into ILVs, where they would also localize to the outer surface of EVs. To investigate these 3 possibilities, we used both siRNA-mediated knockdown and chemical modulators (drugs) targeting relevant pathways. Drug concentrations were selected according to published observations (see Table S2), which we then validated to make sure that the selected concentrations did not impact significantly on cell morphology and viability (Fig. S16).

Knockdown of the ESCRT components *Alix* or *Tsg101* led to a reduction in H3 secretion (Fig. 5A). Annexin A1 was expressed at a low level in OLN-93 cells (Fig. S10B) and EVs (Fig. 1E) and did not change dramatically following siRNA treatments (Fig. 5A), indicating that the microvesicle pathway was unable to compensate for H3 secretion via exosomes. On the other hand, knockdown of the ceramide pathway using an siRNA directed against *neutral sphingomyelinase* (*Smpd2/nSMNase2*) showed only a slight decrease in H3 secretion, which was not significant. Due to the lack of specific antibody recognizing rat Smpd2 in WB, we could not confirm the efficiency of Smpd2 knockdown. Hence, we used the nSMNase2 inhibitor GW4869 to block ceramide production^76, 77^. Treatment of OLN-93 cells with 5 µM GW4869 completely blocked H3 secretion (Fig. 5B).

**Figure 5.**
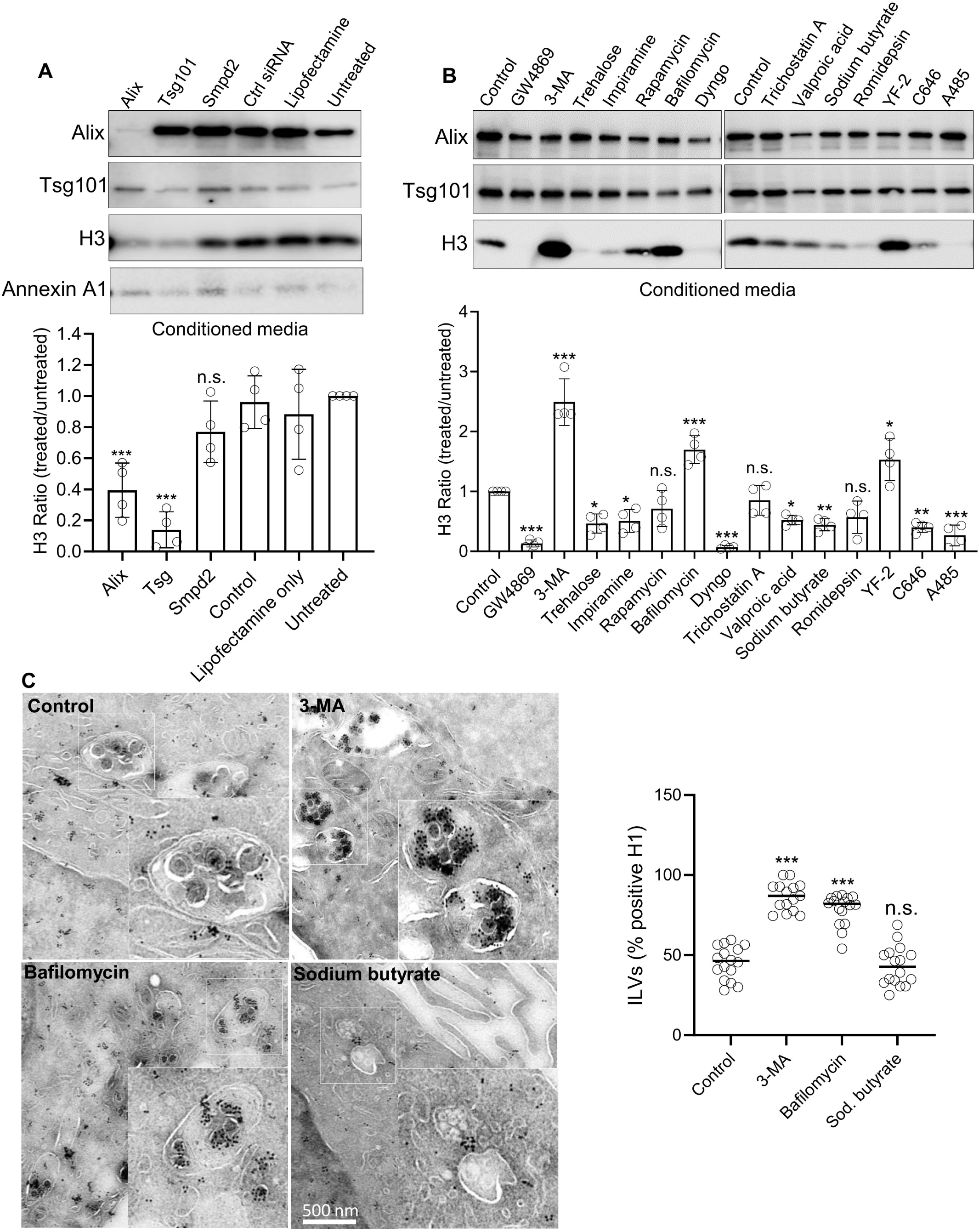
Histone secretion is influenced by multiple cellular pathways. **(A)** WBs showing the effects of siRNA mediated knock-down of *Alix*, *Tsg101*, and *Smpd2* in OLN-93 cells. H3 and EV markers (Alix, Tsg101 as well as the plasma membrane/microvesicle marker Annexin A1) were quantified in total protein recovered from the media by StrataClean pulldown. The experiment was repeated 4 times, and a representative blot is shown. Band intensities were quantified, and the column chart below shows the ratio of H3 (treated/control) from four independent experiments. **(B)** Pharmacological treatment of OLN-93 cells, analysed as in (A). Control is without treatment. GW4869 reduces EV production by the cells, To compensate and visualise Alix and Tsg101 signals in blots 2.5 times more samples were loaded in GW4869 treated samples in comparison to control and other samples. **(C)** Immuno-TEM micrographs showing anti-H1 labelling associated with ILVs in OLN-93 cell sections, following treatment with compounds that significantly increased (3-MA or bafilomycin) or decreased (sodium butyrate) H3 secretion into the media. Right, scatter plot showing quantification of histone positive ILVs under control and following treatment. One-way ANOVA: *, ≤ 0.05, **, *p* ≤ 0.01, ***, *p* ≤ 0.001.

Based on the proteomic analysis of EVs and the non-vesicular secretome, it has been proposed that EV-associated histones may originate from the nucleus and be secreted via an autophagy-dependent amphisomal pathway^33^. To investigate the role of autophagy in histone secretion, we treated OLN-93 cells with the autophagy inhibitor 3-methyadenine (3-MA), which blocks phagophore formation. Treatment with 3-MA increased H3 secretion significantly (Fig. 5B) and a significant increase in histone localization to ILVs (Fig. 5C). On the other hand, inducers of mTOR-independent autophagy (trehalose and imipramine) showed a significant decrease an opposite effect (Fig. 5B). The mTOR-dependent inhibitor rapamycin showed a decrease in H3 secretion that was not significant. Further, we used Bafilomycin A1 (Baf) to block the vacuolar H+-ATPase (V-ATPase) and thereby the fusion of autophagosomes and late endosomes to lysosomes^78^. Like 3-MA, Baf treatment also increased H3 secretion (Fig. 5B) and histone localization to ILVs (Fig. 5C). To investigate the requirement for clathrin-dependent endocytosis, we used the dynamin inhibitor Hydroxy Dynasore (Dyngo-4a)^79^, which downregulated H3 secretion (Fig. 5B).

Histone translocation to, and function in, the nucleus is modulated by post-translational modifications (PTMs), such as acetylation and methylation^80, 81^. Histone acetyl transferases (HATs) modify histones on Lys (L) and Arg (K) residues, thereby reducing the net positive charge of the protein. Histone acetylation is counteracted by the activity of deacetylases (HDACs). To determine if acetylation status is important for histone secretion, we used drugs to either inhibit or activate HDACs or HATs. The pan HDAC inhibitor Trichostatin A as well as Romidepsin (HDAC1 and 2 selective) did not have a significant effect but both valproic acid and sodium butyrate (HDAC1 selective) reduced H3 secretion (Fig. 5B). The p300/CBP selective HAT inhibitors C646 and A-485 reduced, whereas the broad-spectrum HAT activator YF-2 increased, H3 secretion. Hence, blocking ESCRT, ceramide and dynamin pathways inhibited H3 secretion. In contrast, inhibition of autophagosome formation and lysosomal degradation promoted accumulation of H3 in ILV/MVBs and secretion via EVs. Interfering with histone acetylation status modulated H3 secretion though the relationship to acetylation status was unclear.

### Surface association of extracellular vesicle histones and their modifications

During the multiple centrifugation and washing steps used in purification, a significant amount of EV surface proteins may be lost from the corona^82^. Alternatively, non-nuclear histones might fuse with EV membranes under high centrifugal force^83, 84^. Therefore, to confirm that histones are EV surface associated proteins, we used imaging flow cytometry (IFCM) to characterize histone positive EVs in conditioned media without purification^61, 62, 85^. We identified fluorescently labelled anti-human H3 and H4 antibodies with good signal and low noise (Fig. S17A) and combined these with previously characterized anti-human tetraspanin antibodies (CD9, CD63 and CD81). We analysed EVs in the conditioned media of HeLa cell cultures under either control conditions, or following acute exposure to H_2_O_2_(Fig. 6A, S17).

**Figure 6:**
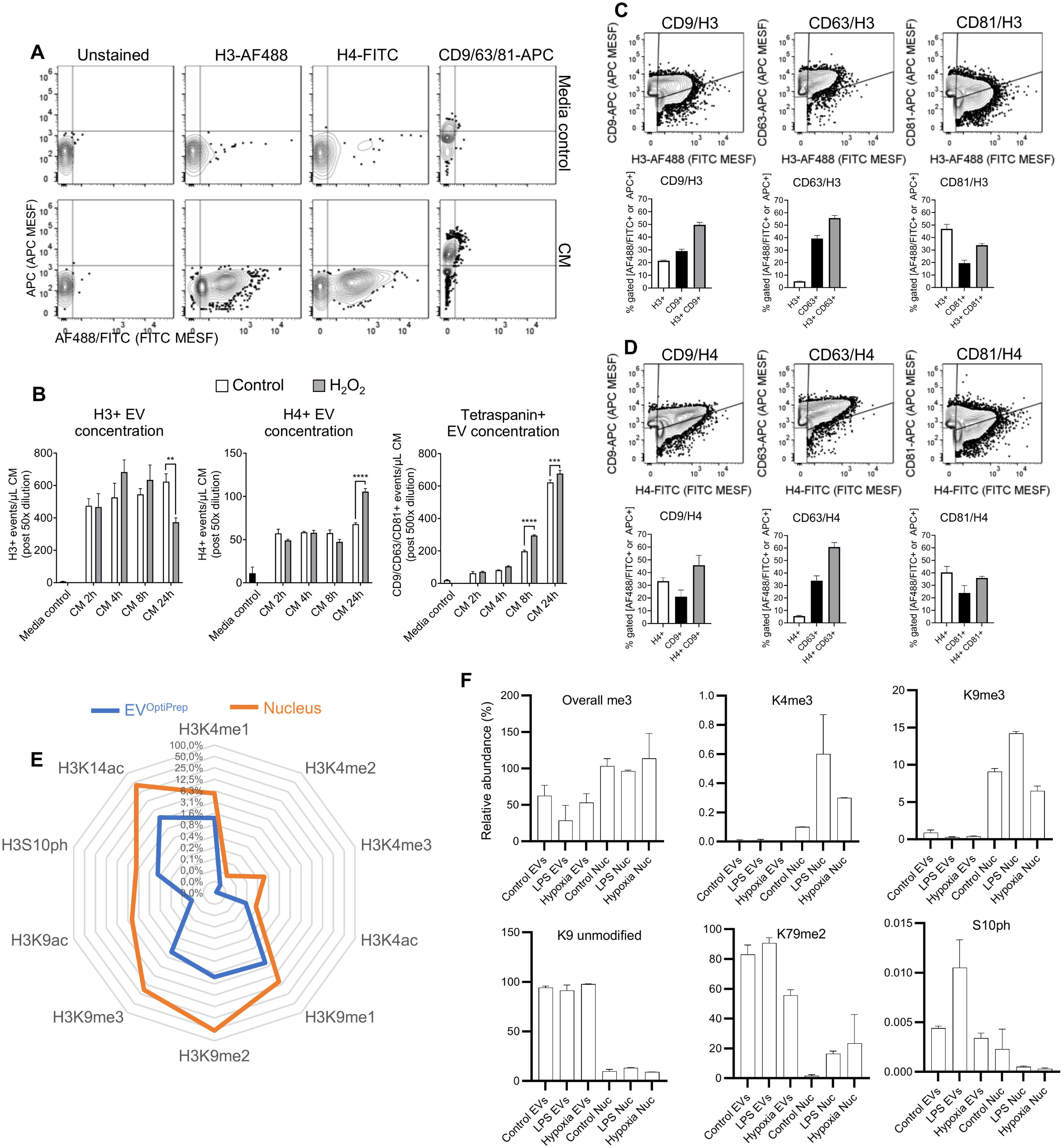
Histones are surface associated in intact EVs and carry less PTMs than their nuclear counterparts. **(A)** Plots showing representative IFCM data obtained from HeLa cell conditioned media (CM) and media controls stained with either AF488 labelled anti-H3 or FITC labelled anti-H4 antibodies, or a mixture of APC labelled anti-CD9/CD63/CD81 antibodies to detect total tetraspanin positive EVs. **(B)** Quantification of detected gateable events positive for single stained H3, H4, or tetraspanins (n=3) in CM samples collected at different time points from both control and H_2_O_2_ treated cell cultures. Samples were diluted post staining as indicated on the *y*-axes. **(C)** H3 co-labelling and **(D)** H4 co-labelling with anti-CD9, -CD63, or -CD81 antibodies in CM from 24 h H_2_O_2_ treated samples. See also Fig. S15. Statistical comparisons by two-way ANOVA. **, ≤ 0.005, ***, *p* ≤ 0.001, ****, *p* ≤ 0.0001. **(E)** Radial plot showing relative abundance of H3 post-translational modifications (PTMs) identified in EV^OptiPrep^ versus nuclear histones. K = lysine, ac = acetylation, me = mono-(me1), di-(me2) or tri-(me3) methylation, ph = phosphorylation. **(F)** Representative examples of H3 PTMs that were selectively enriched in the nucleus, or in EVs. Lysine (K) trimethylation (me3) results from the covalent attachment of three me groups to a K residue and is therefore a relatively “slow” modification. H3 is poorly modified with me3 in EVs relative to nuclei regardless of whether this PTM benchmarks actively transcribed chromatin (H3K4me3) or silenced heterochromatin (H3K9me3). H3K9 is relatively unmodified in EVs, and H3K79me2 is an example of a modification that is enriched in EV histones. Phosphorylation of H3S10 is selectively enriched in EVs from LPS treated cells.

Time course measurements over a 24 h period revealed that the mean fluorescence intensity (MFI) of anti-H3 and -H4 labelling was already maximal by 2 h and remained constant out to 24 h (Fig. S17B). In contrast the MFI of anti-tetraspanin labelling increased from 2-24 h, suggesting that H3/H4 localize to EVs but that the total amount of antibody accessible histone remains constant over time. Interestingly, H3 and H4 showed different kinetics.

Under control conditions, H3 positive EVs showed a gradual increase, with the highest concentration at 24 h. In H_2_O_2_ treated cultures, H3 positive EVs accumulated faster, with highest concentrations measured at 4 h, followed by a gradual decrease (Fig. 6B). Thus, H_2_O_2_ treatment appeared to promote the release of H3 positive EVs. The different kinetics of stress responsive compared to control EVs seen using IFCM may be due to a lower level of antibody accessible H3 on the EV surface. In contrast, H4 positive EVs were detected in control and H_2_O_2_ treated cells at similar levels up to the 8 h timepoint, and their concentration increased markedly at 24 h in the H_2_O_2_ treated cultures (Fig. 6B). This indicates a difference in the kinetics and/or accessibility of antibodies to H3 or H4 on EVs.

To further characterize H3 and H4 positive EVs, we performed double labelling to assess co-occurrence of H3 or H4 together with CD9, CD63, or CD81, on the same EV. About 50-60% of total fluorescence events were found to be double positive for both H3 and CD9 or CD63, while only about one third were double positive for H3 and CD81 (Fig.6C and Fig. S17C). Similarly, H4 was detected more on CD63 positive (∼60%) and CD9 positive (∼45%) EVs than on CD81 positive EVs (∼35%; Fig. 6D and S17C).

To address whether histone PTMs might influence their association with EVs, we characterized the extent and pattern of PTMs on EV associated histones. We extracted histones from OLN-93 EV^OptiPrep^ and the corresponding nuclei from control cells and those exposed to either LPS or hypoxia (Fig. S18) and analysed PTMs using nano-liquid chromatography coupled to high resolution mass spectrometry. Interestingly, common PTMs including acetylation, methylation and phosphorylation were significantly reduced in EV compared to nuclear histones (Fig. 6E). For example, in control EVs single H3K9ac and H3K14ac marks were significantly lower in comparison to the nuclear fraction. More complex modifications, such as trimethylation (me3) of lysine residues in H3 (Fig. 6E, F) were also less abundant in EVs. Hence, histones lacking common modifications were more enriched in EVs, as well as selected mono- or di-methylations (e.g. H3K79me2; Fig. 6F). Phosphorylation on histone H3 (H3S10ph; Fig. 6F), was also more abundant in EVs, particularly following LPS treatment, indicating that a proportion of EV-associated histones may originate from mitotic chromatin. In support of our results with HDAC inhibition or HAT activation, enrichment of unmodified histones in EVs is a novel finding.

## DISCUSSION

Histones are identified consistently amongst the most enriched proteins in EV proteomes^32, 86, 87^. The prevailing view is that EV histones are present in nucleosomes directly associated^30, 31^, or copurifying, with EVs^33, 34^. Alternatively, since histones are such abundant cellular proteins, they may represent a component of the Contaminant Repository for Affinity Purification ome (CRAPome)^49^. It is important to understand both the location and form of EV histones since extracellular histones, cell-free DNA and nucleosomes have different functions^88^. To our knowledge, this is the first study to investigate EV histones without starting from the *a priori* point of view that they are nucleosomes.

Using a range of mammalian cell lines and experimental approaches, including density gradient purification, analysis of ILVs within MVBs prior their release as exosomes, biophysical/chemical fractionation, and IFCM to quantify histones on native EVs, we have found that: i) Histones localize to ILVs before their secretion as exosomes and colocalize on the surface of tetraspanin positive EVs (CD63>CD9>CD81). ii) Histone levels decrease in the cytoplasm and accumulate in MVBs and on EVs in response to cellular stress. iii) The majority of EV histones are bound to the EV membrane and cannot be removed by neutralization of electrostatic interactions. Furthermore, unmodified histones and histone octamers bind efficiently and directly to liposomes and EVs, whereas nucleosomes do not and association of the majority of histone with EVs seems not to require DNA. iv) Histone secretion can be blocked by interference with ESCRT-dependent or independent pathways of EV biogenesis and depend on the autophagosome-lysosome pathway. v) EV-associated histones are distinct from nuclear histones due to their relative lack of PTMs. The possible pathways by which histones may associate with EVs are summarised in Fig. 7.

**Figure 7:**
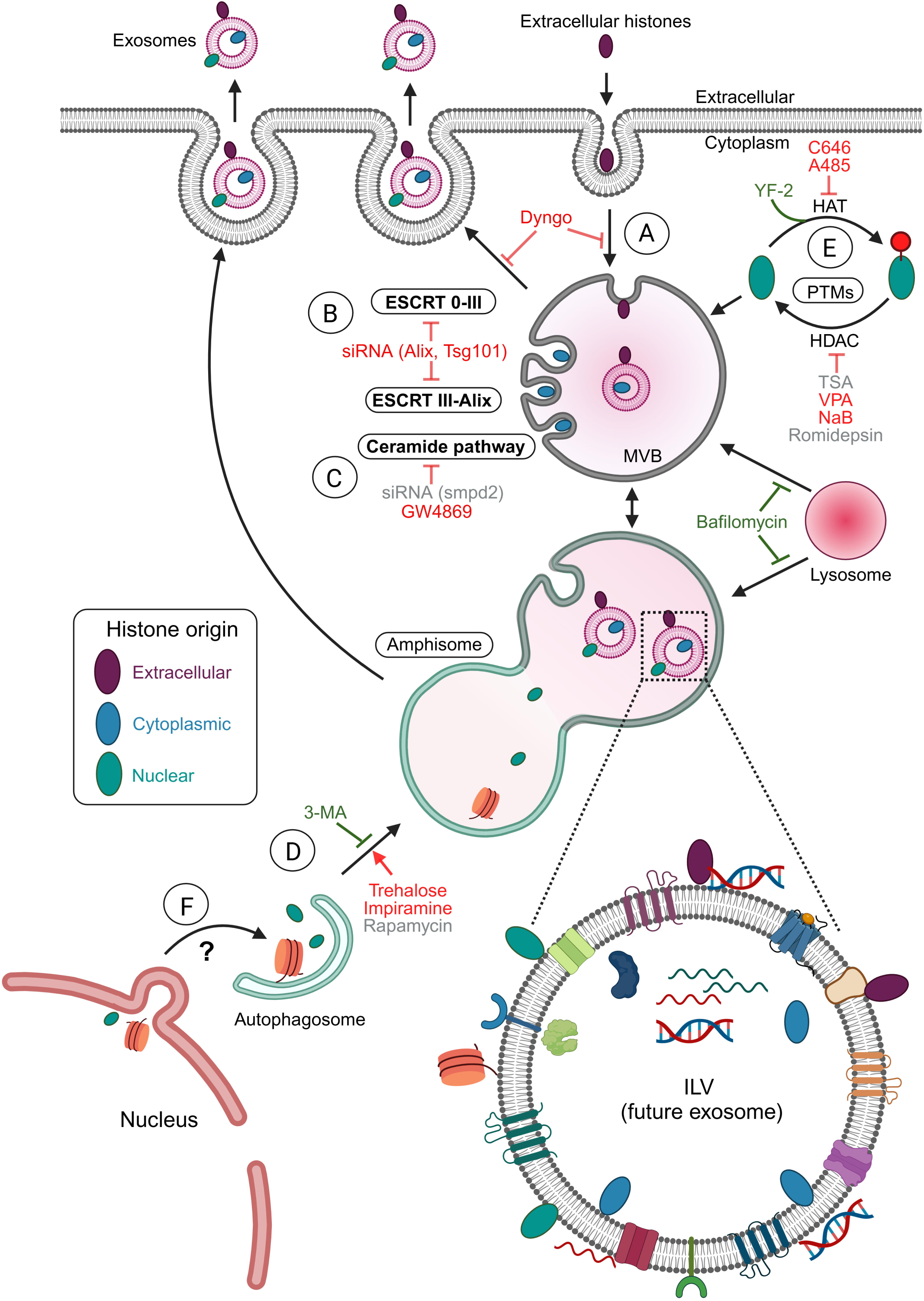
Potential pathways for histone secretion via exosomes. Schematic diagram summarizing the findings of this paper. **(A)** Extracellular histones (yellow diamonds) internalized by endocytosis would undergo inward budding into MVBs and associate with the outer surface of ILVs. Cytoplasmic histones (red diamonds) sorted during **(B)** ESCRT-dependent or **(C)** –independent (ceramide pathway) ILV formation may localize as internal/luminal cargo (red diamonds). **(D)** Histones originating from the nucleus (green diamonds) and captured by autophagosomes may partition to ILVs after amphisomes or lysosomes fuse with MVBs. **(E)** Posttranslational modification of histones in subcellular compartments such as the cytoplasm may also be important for sorting to MVBs/ILVs. **(F)** The possible route whereby histones may traffic from the nucleus to amphisomes is unknown.

Histones are such well characterized chromatin proteins it is natural to assume that EV histones are also DNA associated and exist as nucleosomes. A seminal study identifying histones in dendritic cell derived exosomes^34^ noted core histone association with apoptotic material with a relatively high density (1.16-1.28 g/ml), which would equate to fractions 10-12 on the OptiPrep gradients used in this study. Our findings indicate that whilst histones localize to higher density fractions 10-12 in EVs from certain cell lines (e.g. AC-16), or following cell stress in OLN-93 cells, this is the exception rather than the rule. The findings of Théry *et al.* then contributed to a leading review that proposed histones as luminal cargo within large microvesicles and/or apoptotic vesicles, rather than being on the surface of exosomes^43^. More recently, it was proposed that histones are secreted as a complex with dsDNA and can be separated from EVs in the non-vesicular fraction^33^. Our findings do not support this hypothesis. The enrichment of chromosomal DNA in large microvesicles has been confirmed^89^. However, such large EVs would have been removed by the purification method used here. Other studies have addressed the localization of DNA at the surface of EVs^30, 31^, though neither of these have investigated the direct binding of free histones to EVs. Importantly, one factor that has been overlooked consistently is the lack of stoichiometry reported for core histones detected in EV proteomes^32^. As a specific example, the expected chromatin/nucleosome H3:H4 ratio of 1:1 is detected in quantitative proteomics of cell lines by Super-SILAC^90^. However, the same stoichiometry of core histones is not a consistent feature of in EVs^86^, which is confirmed in non-SILAC proteomics of highly purified EVs intended for therapeutic application^91^. Hence, a parsimonious explanation of the available data is that although nucleosomes may associate with EVs under certain circumstances and/or cell types, most EV histones are not nucleosomes. Our results suggest that non-nucleosome histones comprise a significant component of sEV histones and that they are not dependent on DNA for EV association. This calls for a careful re-evaluation of the form of histones and DNA present in EVs subpopulations from different cell types and we provide methodological approaches to achieve this.

In agreement with proteomic analysis of EV membrane proteins^92^, we found that only a small fraction of EV-associated H3 is found in the luminal fraction of OLN-93 EVs. The majority of H3 was membrane associated and we identified both a weak interaction with the EV surface (disrupted with 750 mM NaCl), and a much stronger interaction with the membrane fraction that could not be disrupted by ionic interference. Could histone EV binding be mediated primarily by by be due to protein-protein or. protein-glycan interaction? In *Drosophila*, H2A and H2B are associated intracellularly with lipid drops^93^, which function as a sink to supply histones for cell division and as antimicrobial effectors during bacterial infection^94, 95^. Unilamellar lipid drop binding is mediated by the protein chaperone Jabba^95, 96^ and although histone-lipid drop association is conserved in mammals^94^, a functional homologue for Jabba has not been identified. We have not investigated specifically whether histone-protein interaction contributes to EV binding, but our data shows that all histones can bind independently to purified membrane derived liposomes. These findings agree with previous reports. Histones bind directly to membrane phospholipids^97^ and binding may occur via the phosphodiester bonds of phosphate groups within phospholipids, mimicking the binding of histones in nucleosomes to the phosphodiester bonds in DNA^12^. This is an important point as it predicts direct competition between phospholipids and DNA for histone binding, though this remains to be tested

Interestingly, there is also evidence for membrane penetration^97^ and translocation^98, 99^ of histones. Penetration was enhanced by negative charge and inhibited by cholesterol, indicating that phospholipid content and membrane asymmetry is important for histone binding. In native cell membranes or EVs, negatively charged phospholipids are concentrated at the inner leaflet, with positively charged phospholipids at the outer surface^100^. In liposomes, this membrane asymmetry is lost, but in EVs it is unclear how histones might gain access to negatively charged phospholipids on the inner leaflet. Since the negatively charged phosphatidylinositol (PI) derivative PI(3,5)P_2_ is enriched in endosomes-MVBs-lysosomes^101^, it is tempting to speculate that PIP_2_ plays a significant role in histone recruitment and sequestration into ILVs/exosomes. Therefore, it is important to determine the affinity of individual and multimerized histones, with and without DNA, for lipid membranes with different phospholipid composition to see how the lipid environment dictates histone binding capacity and location/conformation.

Upregulation of histone secretion was a common response to cellular stress. However, interesting differences were observed in the kinetics, amount, and floating density of histone positive EV fractions. Upregulation of EV histones was observed following heat stress, together with a smaller EV size. In contrast, the levels of cytoplasmic histones were reduced but within MVBs, histones increased, suggesting that stress induces the sequestration and secretion of cytoplasmically located histones. LPS challenge has also been shown to lead to an upregulation of EV-associated histones in mouse bone marrow-derived macrophages, which was proposed to be due to migration of nuclear histones to the cytoplasm^102^. Whatever the source of cytoplasmically located histones, accumulation can activate signalling pathways that may lead to apoptosis^19, 103^. Hence, cells may sequester and secrete histones as part of a stress-associated survival mechanism, whereby vesicles are released continually but histones are loaded into them in a regulated manner.

In clinical contexts, circulating extracellular histones have been found in various pathological^19, 104, 105^ as well as non-pathological^106^ states. Our findings support the idea that at least some of these circulating histones could be EV associated. However, it is unknown whether EV associated histones have a different function compared to other types of extracellular histones identified in sepsis, trauma, or cell death^19, 23^. EVs might represent a mechanism by which excess histones can be distributed to other cells^99, 107^. Alternatively, histones may populate the secretory and endosomal pathways and function as innate immune factors to rapidly respond to pathogenic stimuli, whilst buffering the detrimental effects of their highly cationic sequences. Though our findings do not address these questions, they do highlight the importance of accurate characterization of both the nature and the origin of extracellular histone/DNA complexes since they may have different biological functions as well as pathological effects.

EV associated histones may recycle from the extracellular space via endocytosis, as treatment with Dyngo reduced histone secretion (Fig. 7A). However, we also found that both ESCRT-dependent and ESCRT-independent pathways of EV biogenesis were important for histone secretion (Fig. 7B, C) suggestive of an intracellular source. MVBs are a key node in the regulation of secretory turnover, balancing degradation (via late endosome/lysosome fusion) with recycling (via fusion to the plasma membrane)^108^. Modulation of autophagy and lysosome maturation significantly affected the secretion of histones. Inhibition of phagophore formation (3-MA treatment: Fig. 7D) led to a strong upregulation of H3 in EVs. Activation of autophagy (trehalose, impiramine) had an opposing effect. Furthermore, inhibition of lysosomal degradation (Baf) increased H3 secretion. Hence, this supports the conclusion that a proportion of histones normally degraded via this pathway may be routed to MVBs for secretion via exosomes. This is significant because histones lack a signal sequence and thus sorting into MVBs/ILVs should occur via a non-canonical pathway. Recent findings have begun to define pathways by which cytoplasmic histones are targeted to lysosomes for Lamp 2A-mediated degradation^109, 110^. An amphisome/lysosome origin for EV histones could also explain why most histones associate with the outer surface of EVs, rather than the lumen, as would be the case if they were sorted through canonical ESCRT-dependent-or ceramide pathways from the cytoplasm. After being translated and before being incorporated into chromatin, histones lack most modifications apart from the well-documented exception of H3K9me and H4K5ac and K12ac^111–113^. Other modifications are deposited on histones after their incorporation into chromatin. Since our data shows the presence of H3K27, H3K36, and other H3 and H4 methyl PTMs on EV-associated histones, it indicates that these histones likely originate from nuclear chromatin. Hence, although our data points towards a nuclear source for histones and the importance of autophagy, it is unclear how cells may regulate the routing/sorting of histones and/or DNA through MVBs and into ILVs/EVs, highlighting this as an important area for further study.

Our results do suggest that the removal of histone PTMs may be important for the sorting/secretion process. PTMs are known to regulate the localization of histones to^114^ and within^115^ the nucleus. The most studied PTMs are acetylation and methylation that are involved in the regulation of gene expression^116^ and are primarily regulated via HAT and HDAC enzymes. HAT activation via YF-2 increased histone secretion, whereas HAT inhibition showed the opposite effect. Inhibition of HDAC activity also led to significant decreases in EV-associated H3 (Fig. 7E). Reduction of neutral PTMs on K/R residues would increase histone net positive charge and may promote histone binding to lipids during the sorting process. Since, we could not easily determine how modulation of PTM status related to histone secretion, we determined the extent of histone PTMs to understand how they might localize to EVs. The most striking finding was that EV associated histones are relatively unmodified compared to their nuclear counterparts. Marks such as H3K4me3 and H3K9me3 were almost completely absent in EVs. These two PTMs are the most well characterized occurrences of trimethylation and define active gene expression and constitutive heterochromatin, respectively. These findings highlight that it is not just histones from open or closed chromatin that translocate to EVs; we observed a general loss of modifications that are known to have critical regulatory role in the nucleus.

## CONCLUSION

Our results show clearly that membrane associated histone proteins are secreted specifically via the MVB/exosome pathway. Secretion is regulated by cellular stress responses, leading to an upregulation of histones associated with a population of smaller sized EVs. Unexpectedly, secreted histones do not depend on DNA for EV association and lack many of the of PTMs of their nuclear counterparts. Understanding how histones are routed to and sorted in to MVB for secretion, how they bind to membranes, as well as their functional significance are the next steps.

## DATA AVAILABILITY

Mass spectrometry raw data availability. All raw mass spectrometry data files from this study have been submitted to the Chorus repository (https://chorusproject.org/pages/index.html) under project number 1847.

## AUTHOR CONTRIBUTIONS

Conception: SELA, MH, JG. Design: All authors. Execution: BS, MS, UM, BN, AG, StSt, LS, SiSi, JDG. Writing draft: BS, MS, UM, BN, MH, JDG. Reviewing, and editing: BS, MS, MH, StSt, AG, JZN, OPBW, SiSi, SELA, MH, JDG. All authors read and approved the final manuscript.

## ACKNOWLEDGEMENTS

We are most grateful to colleagues who have contributed time and materials and thank Prof. C. Richter-Landsberg (Department of Biology, University of Oldenburg, Germany) for providing OLN-93 cells and Dr. R. Lodge (Department of Medicine, Laval University, Canada) for the pEGFP-C2 CD63-GFP. Hanna Levén May and Iwan Jones generated the HA-H3-Flag construct, Sophie Rome, Giulia Corso and Mattias Hällbrink provided valuable technical advice, and Fazal Oozeer and Adrian Pini contributed with early insights into extracellular histones. Fransesca Aguilo supplied us with AChE and HDAC1 antibodies to verify nuclear/cytoplasmic extracts and provided valuable input into the project. We gratefully acknowledge Chaitali Chakraborty for assistance with TapeStation, the Biochemical Imaging Center at Umeå University and the National Microscopy Infrastructure, NMI (VR-RF 2019-00217) for microscopy support, and Umeå Centre for Electron Microscopy (UCEM) in association with the NMI, VR-RFI2019-00217 and VR-2018-06478 for EM support and method development.

The study was supported by Umeå University, including the Foundation for Medical Research (Insamlingsstiftelsen), Stiftelsen Forskningsfonden För Klinisk Neurovetenskap Vid Norrlands Universitetsskjukhus, Västerbotten County Council/Region Västerbotten, Kempe Foundations, Stratneuro Initiative and Neurofonden.

