## Supplementary figures for "Histones are exosome membrane proteins regulated by cell stress"

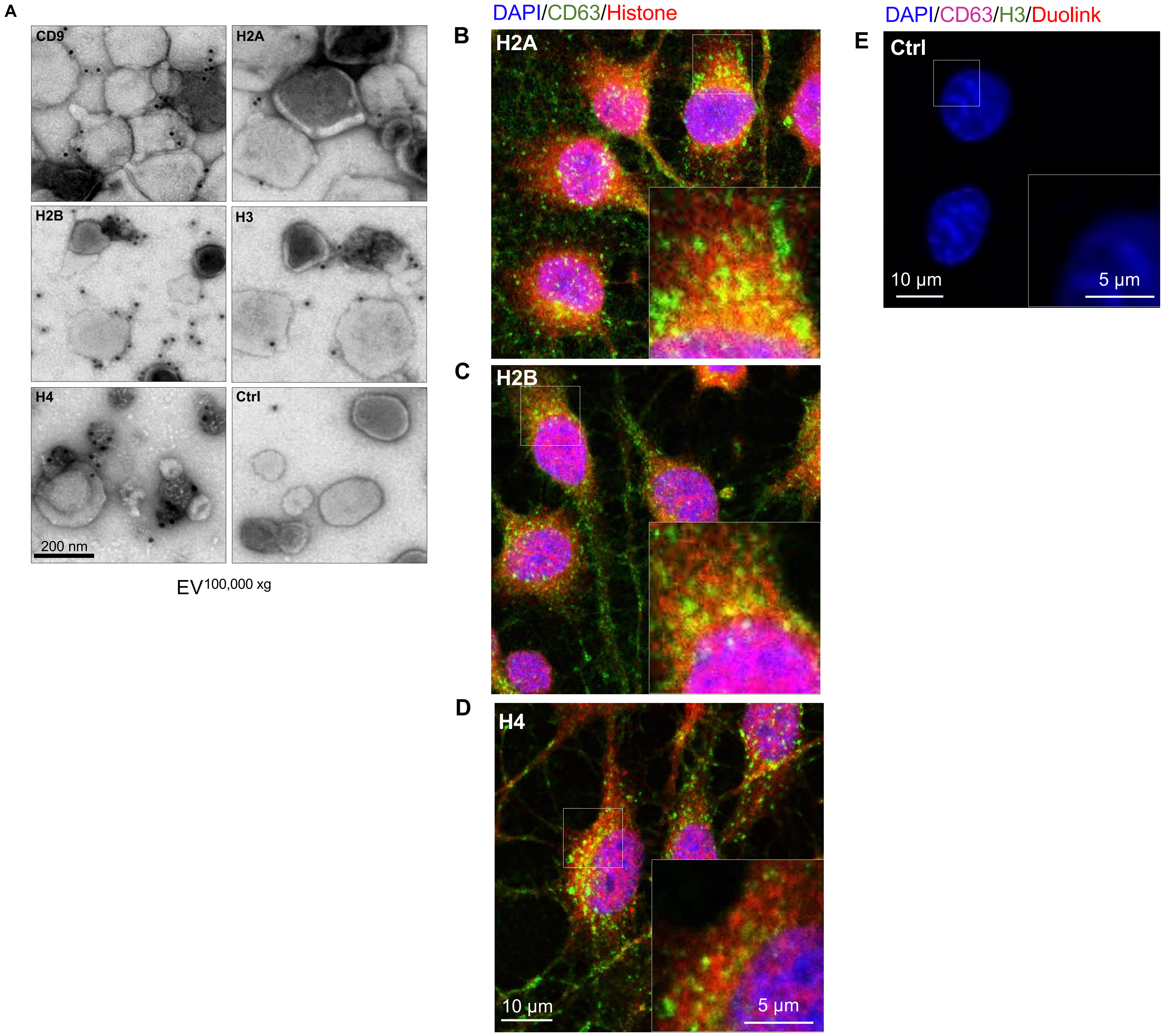


**Supplementary Figure S1. (A)** Immuno-TEM images showing anti-tetraspanin CD9 (EV marker) and anti-histone H2A, H2B, H3 or H4 at the surface of OLN-93 $EV^{100,000\ xg}$. Control (Ctrl) is without primary antibody. Scale bar in H4 = 200 nm. **(B-D)** Confocal micrographs from a single z-plane of OLN-93 cells showing colocalization of anti-histone antibodies (red; as indicated) with anti-CD63 (green). Nuclei were counterstained with DAPI (blue). Inset shows a cropped and expanded image of the white boxed region in the main panel. Scale bar = 10 µm in main panels or 5 µm in insets. **(E)** Confocal micrograph as in (B-D) showing a control sample for (Fig. 1D), Duolink assay performed in the absence of primary antibodies.

A

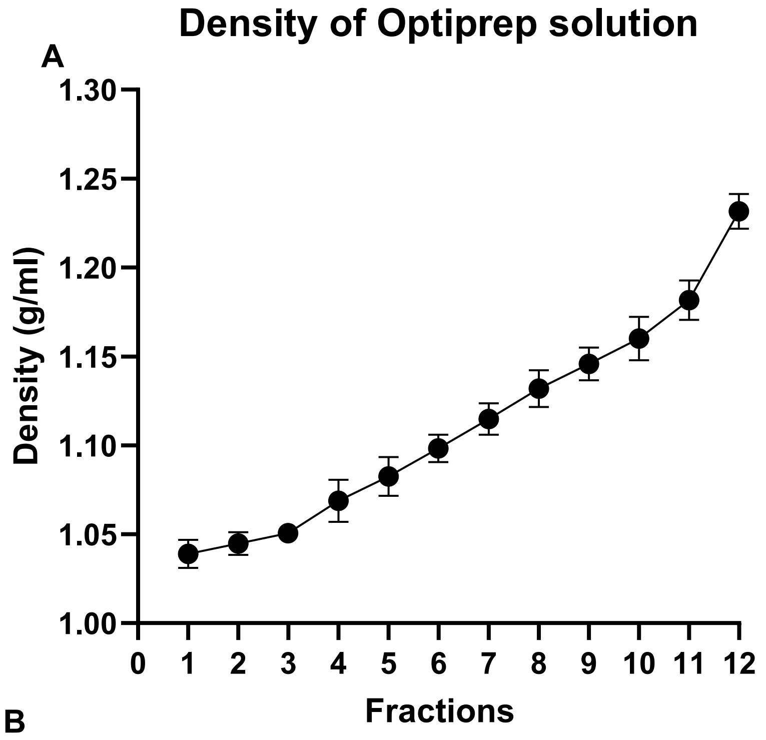


| Fractions | Density (g/ml) |
| --- | --- |
| 1 | 1.03±0.007 |
| 2 | 1.04±0.006 |
| 3 | 1.05±0.005 |
| 4 | 1.06±0.011 |
| 5 | 1.08±0.010 |
| 6 | 1.09±0.007 |
| 7 | 1.11±0.008 |
| 8 | 1.13±0.010 |
| 9 | 1.14±0.009 |
| 10 | 1.16±0.012 |
| 11 | 1.18±0.011 |
| 12 | 1.23±0.009 |

Fractions 4-7 pooled for analysis

B

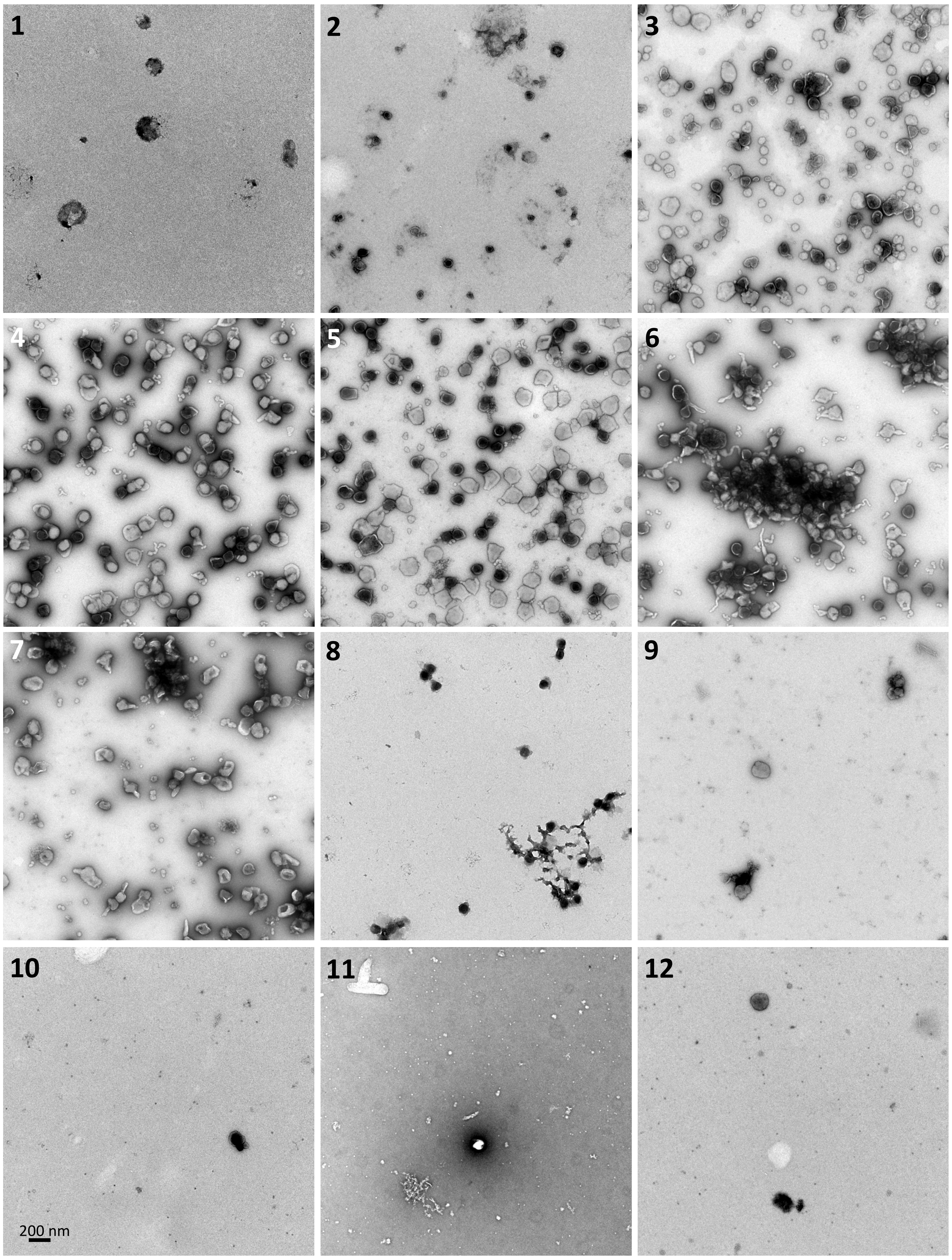


**Supplementary Figure S2. (A)** Line graph showing density of OptiPrep fractions derived from the absorbance (Nanodrop spectrophotometer) at 340 nm using a control gradient (without EVs). Table to the right shows the mean density of each fraction derived from 3 measurements and the SD. **(B)** Representative TEM images of fractions 1 through 12 after separation of OLN-93 EVs$^{100,000\,xg}$ on OptiPrep. EVs localized predominantly to fractions 3-7. Scale bar in 10 = 200 nm.

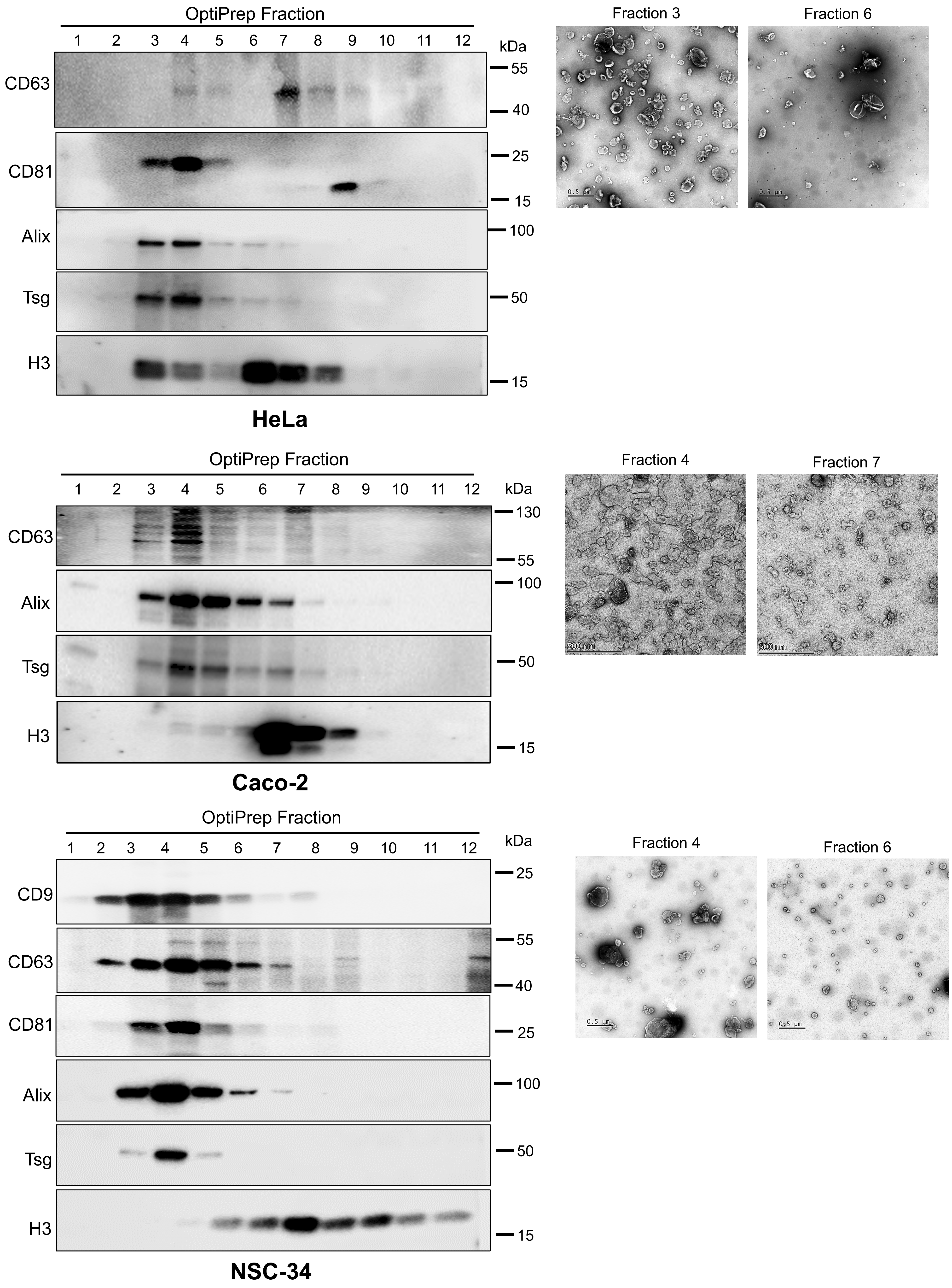


**Supplementary Figure S3. (Left)** Western blots of OptiPrep density gradient fractions of EVs from cell lines, as indicated. Different fractions showing the localization of CD63, CD81, CD9, Alix, Tsg101 (Tsg), and histone H3. **(Right)** Representative TEM images of EV-containing fractions. Scale bars = 0.5 µm/500 nm.

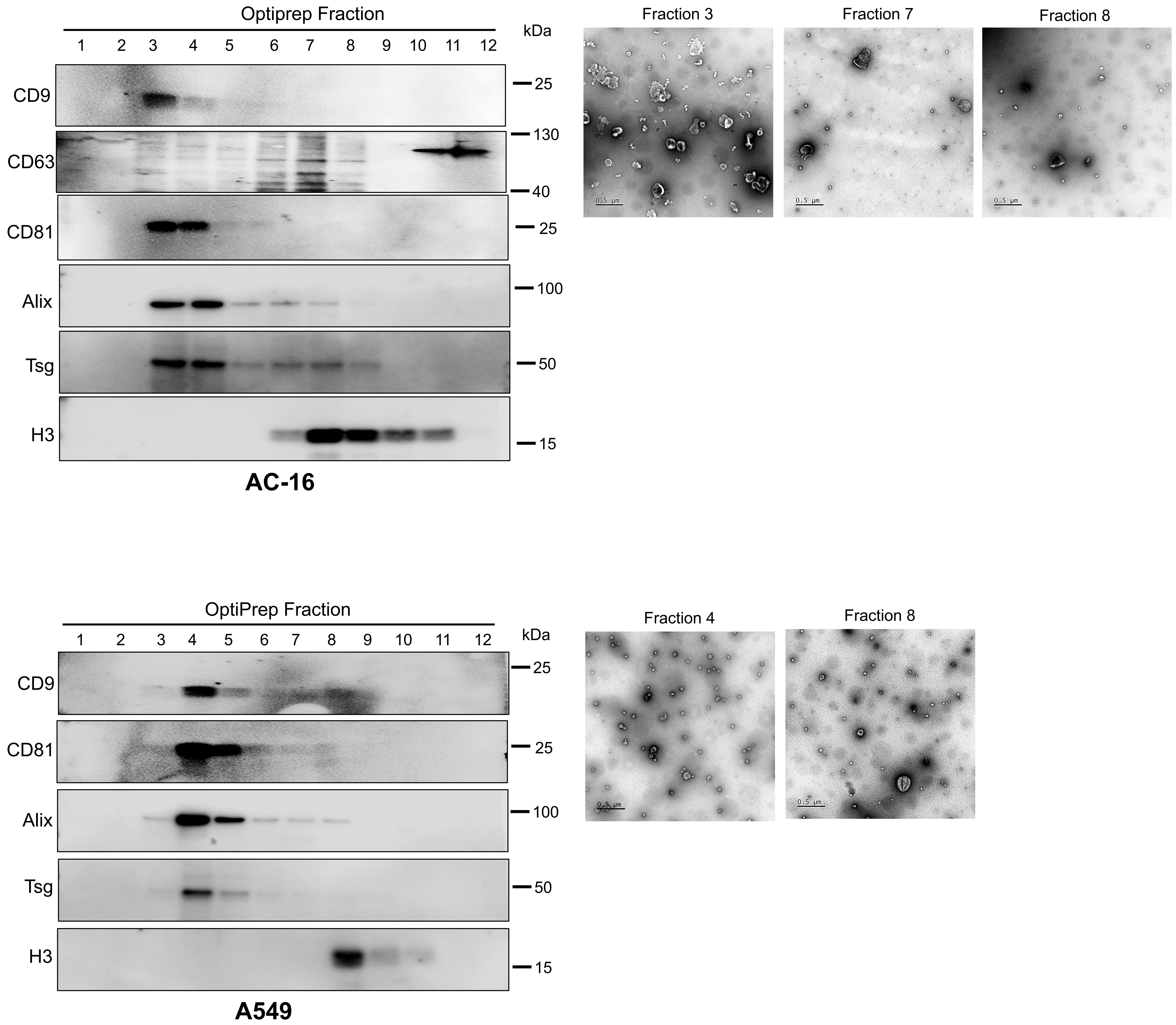


**Supplementary Figure S4. (Left)** Western blots of OptiPrep density gradient fractions of EVs from cell lines, as indicated. Different fractions showing the localization of CD63, CD81, CD9, Alix, Tsg101 (Tsg), and histone H3. **(Right)** Representative TEM images of EV-containing fractions. Scale bars = 0.5 µm/500 nm.

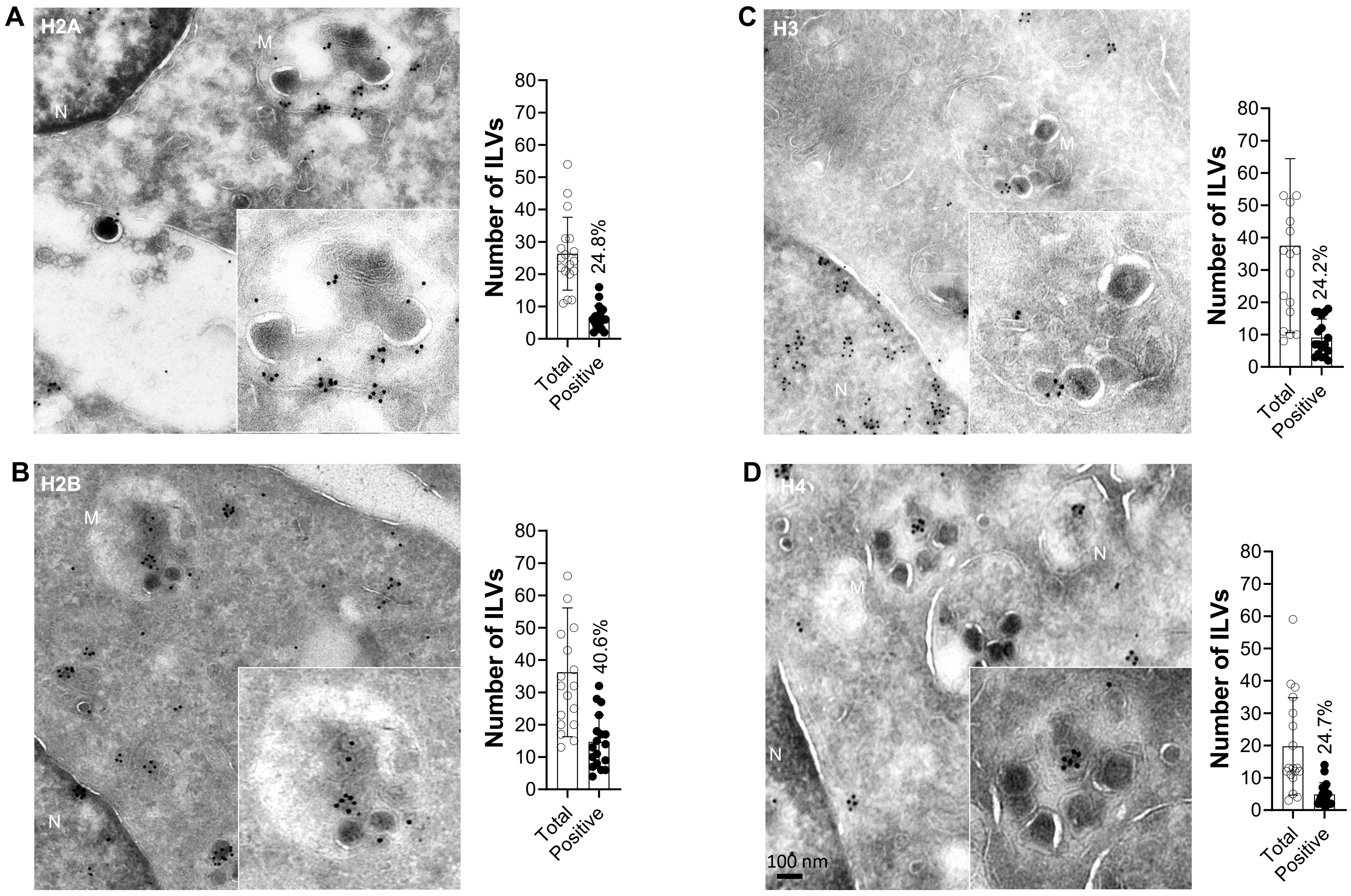


**Supplementary Figure S5. (A-D)** Immuno-TEM on ultrathin sections of OLN-93 cells showing anti-H2A, H2B, H3 and H4 labelling. Insets show a zoomed region of the main image illustrating staining of ILVs within an MVB. A total of 10 images from each grid were analysed for total ILVs and ILVs that contain at least 1 gold particle (considered positive). Column charts to the right of each panel show quantification of total number of ILVs per MVB (left column) and the total number of histone positive ILVs (right column). Percentages refer to % positively labelled ILVs compared to the total ILVs. Error bars represent the mean ±SD. Scale bar in E = 100 nm.

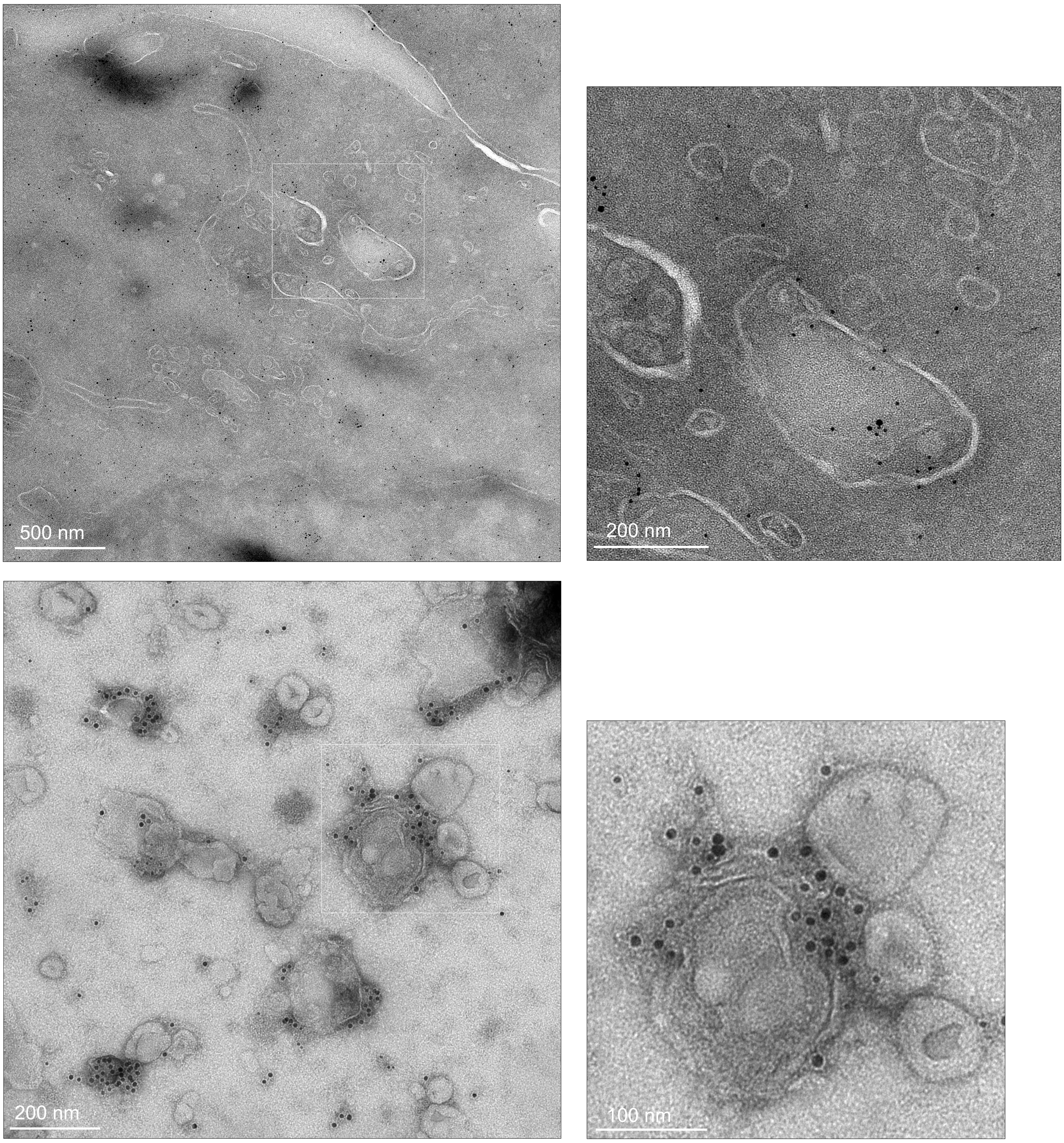


**Supplementary Figure S6. (A)** Immuno-TEM showing a section of a HeLa cell probed with anti-histone H1 (5 nm), and anti-CD63 (10 nm) gold particles. White boxed area shows an MVB, which is magnified in the right image. **(B)** Purified HeLa EV$^{OptiPrep}$ (Fraction 3). White boxed area shows the region of the image shown magnified on the right. Scale bars = 500, 200 or 100 nm as show in each panel.

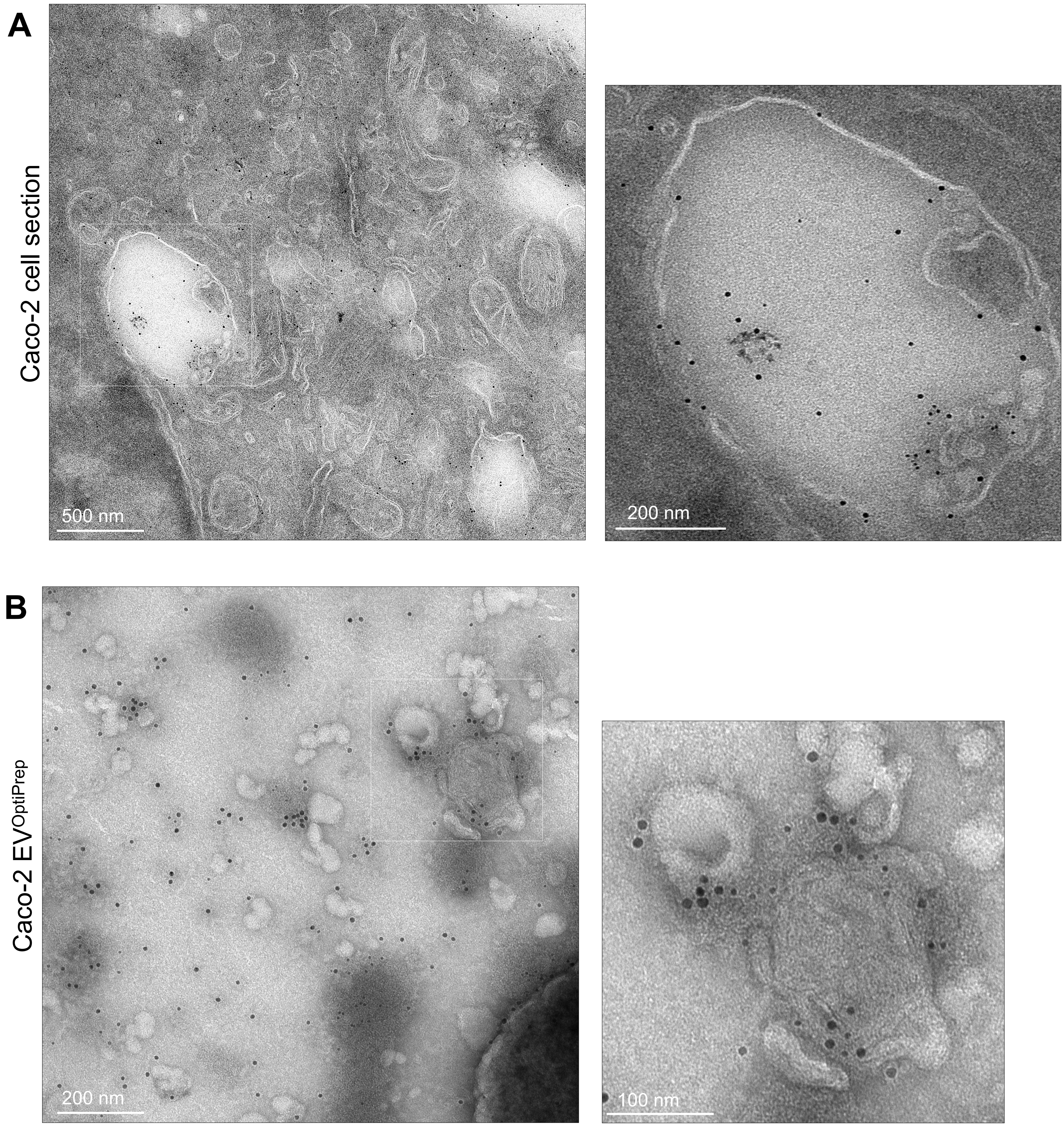


**Supplementary Figure S7**. **(A)** Immuno-TEM showing a section of a Caco-2 cell probed with anti-histone H1 (5 nm), and anti-CD63 (10 nm) gold particles. White boxed area shows an MVB, which is magnified in the right image. **(B)** Purified Caco-2 EV$^{OptiPrep}$ (Fraction 7). White boxed area shows the region of the image shown magnified on the right. Scale bars = 500, 200 or 100 nm as show in each panel.

Figure S8, Singh *et al.*

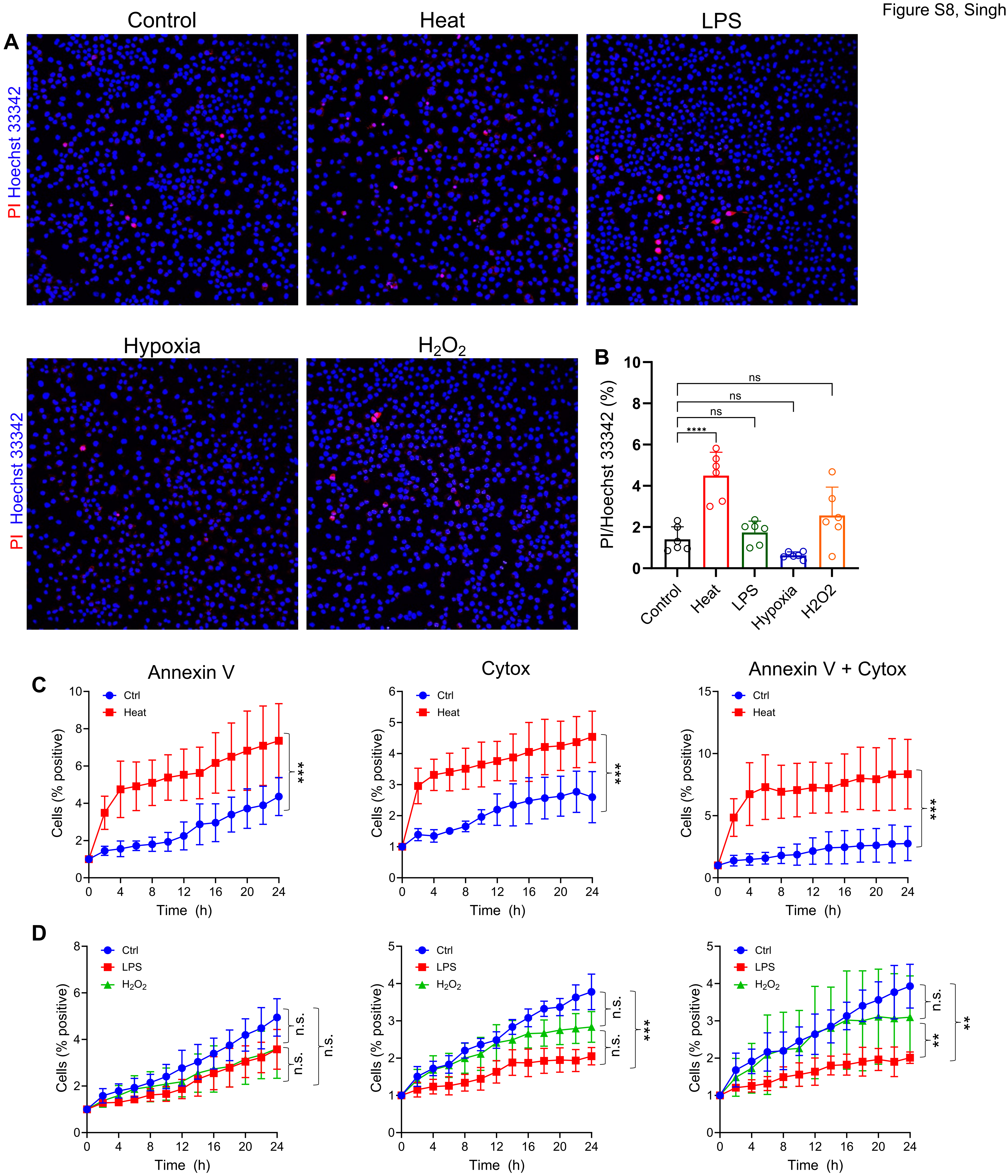


**Supplementary Figure S8.** Cell viability and apoptosis assays of OLN-93. **(A)** Representative fluorescence micrographs after live cell staining with propidium iodide (red; PI, stains dead cells) and Hoechst 33342 (blue; also stains live cells). Significant increase in cell death was observed for Heat but not for other stresses. **(B)** Column chart showing percentage of dead cells (% PI/Hoechst; n = 6 random images). Statistical calculations were performed using Two-way ANOVA and Bonferroni test. **(C-D)** Apoptosis assays performed using Annexin V-red (pre-apoptotic/apoptotic cells), and Cytox-green (dead cells). Double staining indicates cell death by apoptosis. Cultures were labelled and analysed using Incucyte. Significant increase in apoptosis was detected after Heat, but not for $H_2O_2$ and LPS resulted in a significant decrease in apoptosis. For panel C, non-parametric T-test was used, for panel D One-way ANOVA and Dunnet`s test was used. *** p< 0.0005, ** p<0.005.

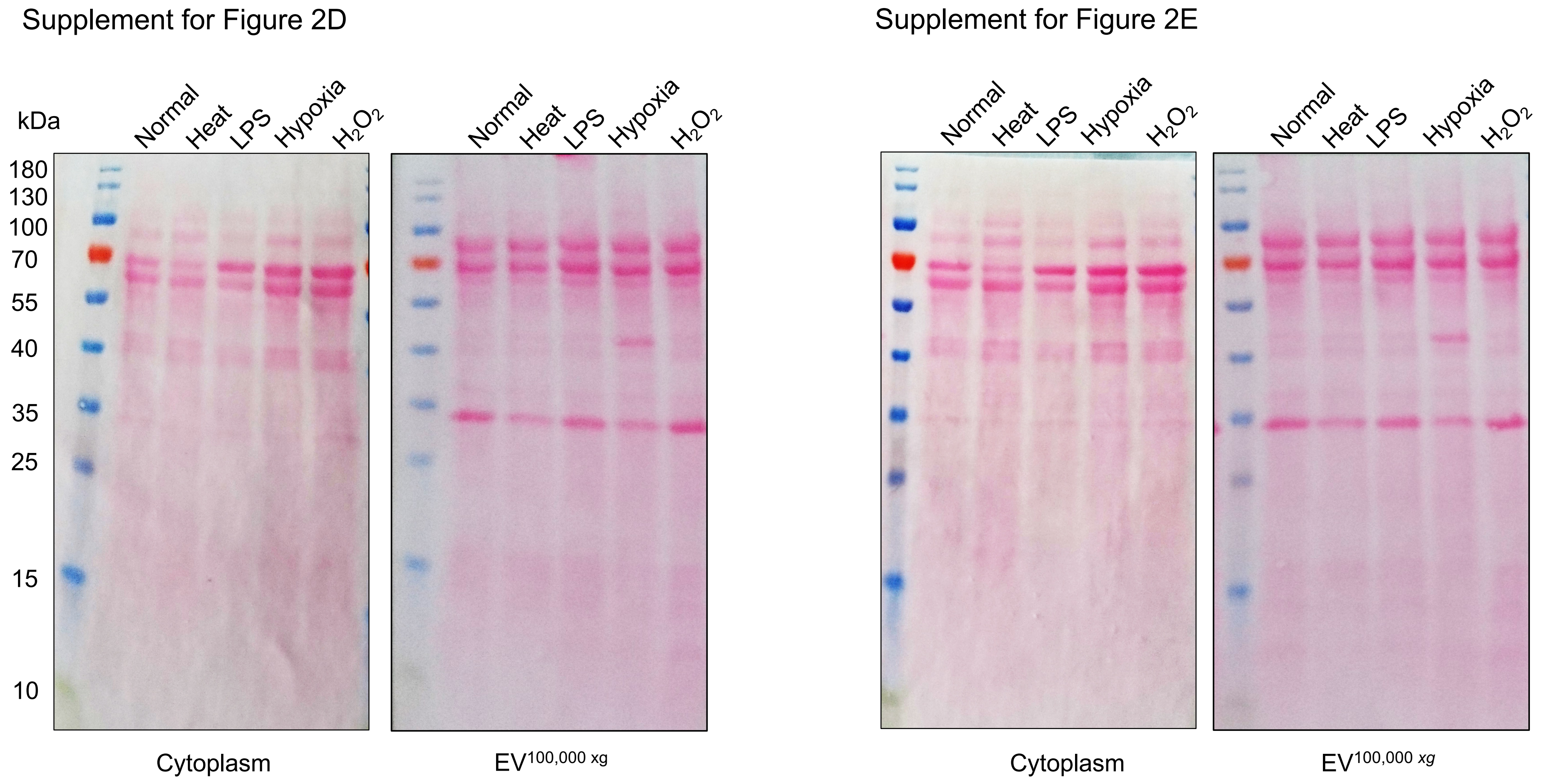


**Supplementary Figure S9.** Ponceau S staining of nitrocellulose membrane blots. Images corresponding to Fig. 2D and Fig. 2E.

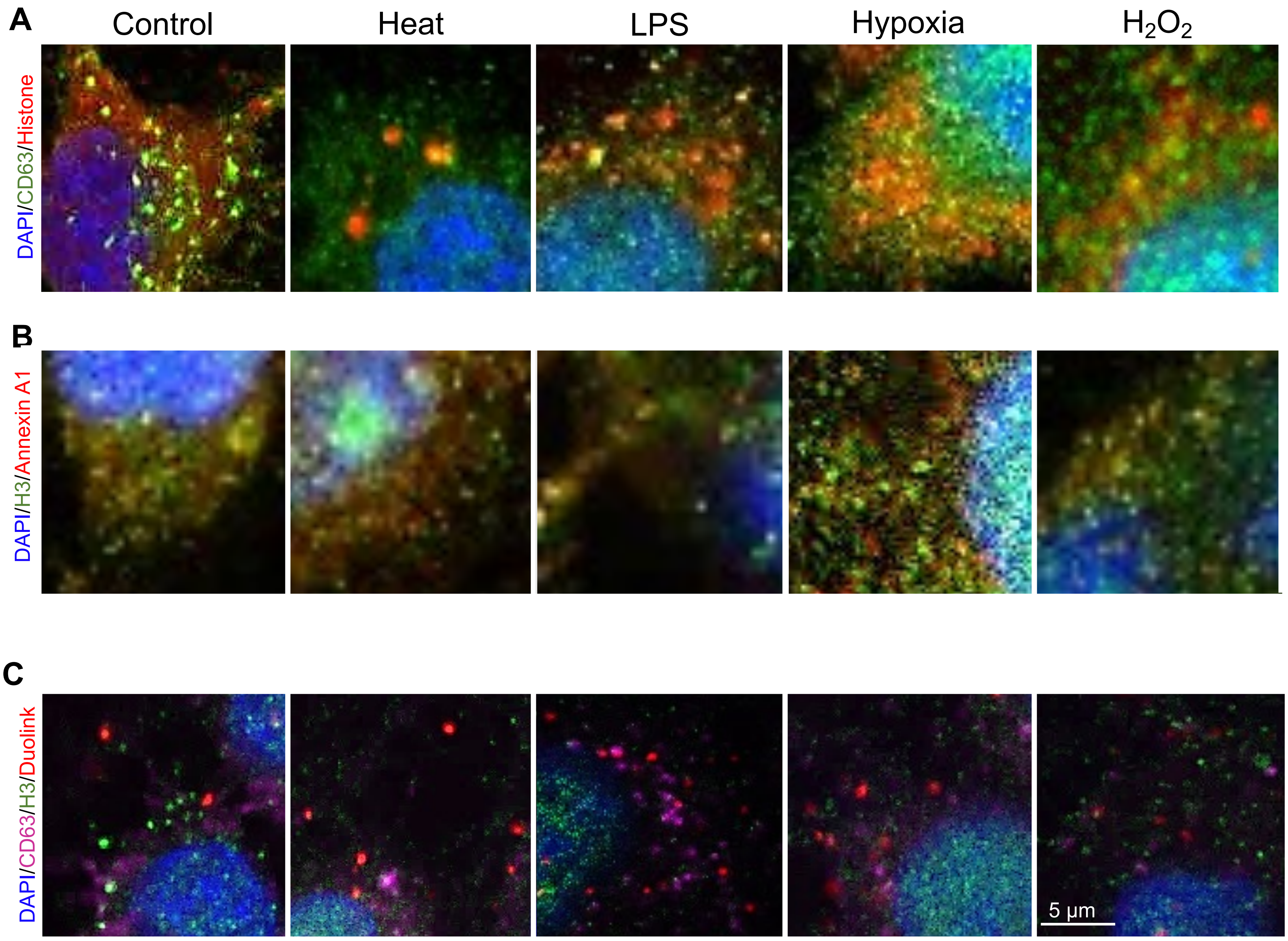


**Supplementary Figure S10. (A)** Confocal micrographs (single z-plane) showing immunofluorescence from OLN-93 cells grown under control conditions or following exposure to different stressors. Anti-CD63 (green) and anti-H3 (red). DNA counter stained with DAPI to label the nucleus (blue). Yellow staining indicates areas of colocalization. **(B)** OLN-93 (control and stress) stained with Anti-H3 (green) and anti-Annexin A1 (red). **(C)** Confocal micrographs of OLN-93 cells additionally subjected to a proximity ligation assay (Duolink; red) between anti-CD63 (mouse; magenta) and anti-H3 (rabbit; green). Red spots indicate proximity labelling between sites of anti-rabbit and anti-mouse immunoreactivity. Scale bar = 5 µm.

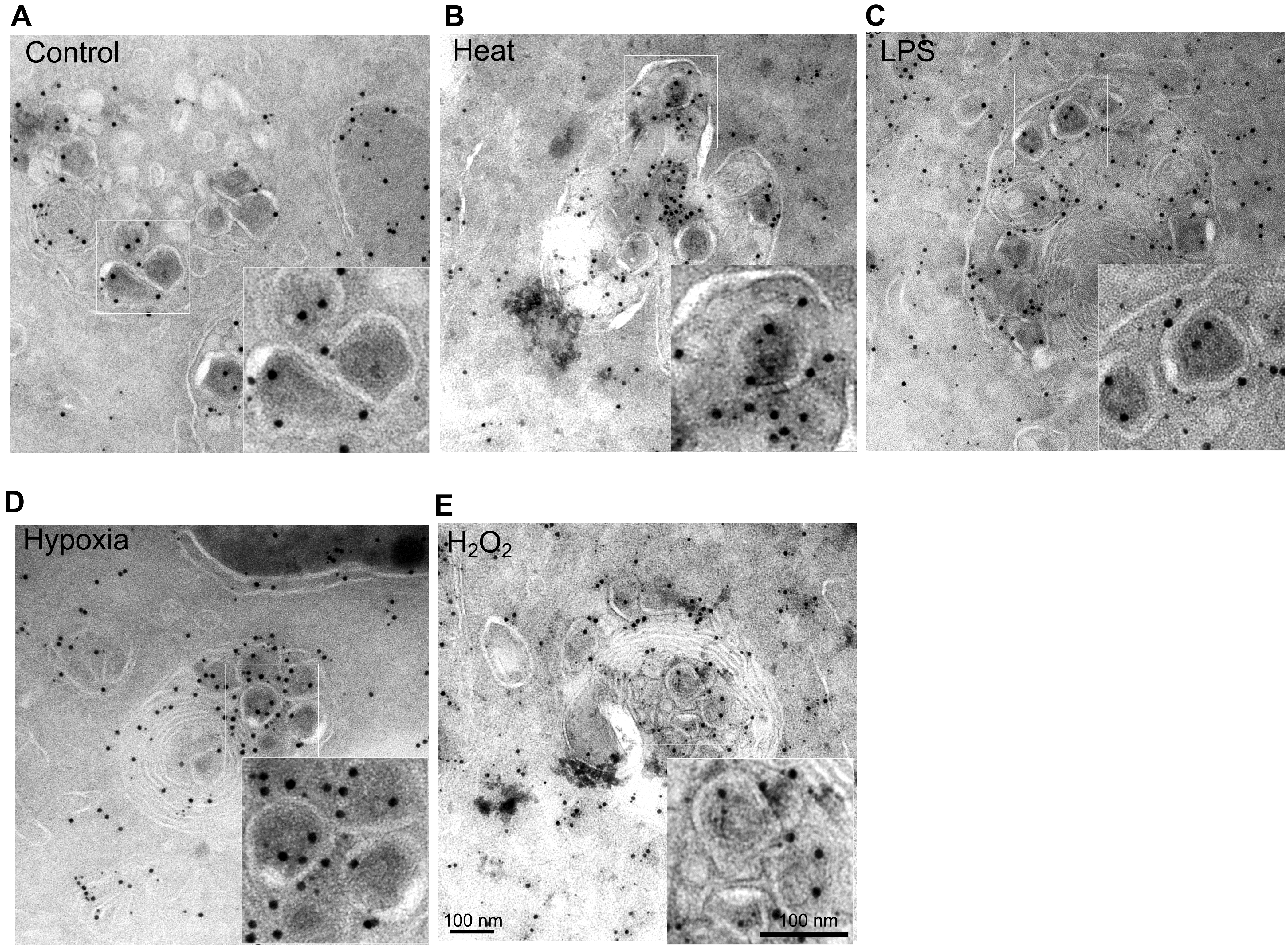


**Supplementary Figure S11. (A-E)** Immuno-TEM of ultrathin sections of OLN-93 cells stained with anti-histone H1 (10 nm gold) and anti-CD63 (5 nm gold). Insets show a higher magnification of an MVB (white boxed region). These are full images of Fig. 2F. Scale bars = 100 nm.

Figure S12, Singh *et al.*

A

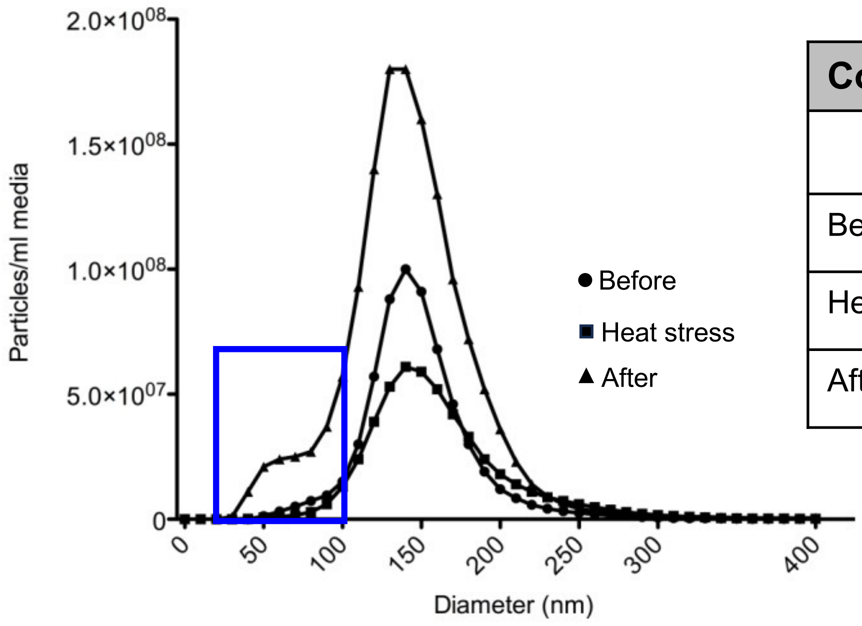


B

| Condition | EVs | | | | | |
|---|---|---|---|---|---|---|
| | **Particle No. (n ml$^{-1}$)** | **Protein (µg ml$^{-1}$)** | **Mean size (nm)** | **Mode size (nm)** | **Conditioning period (h)** | **Rate (n ml$^{-1}$ h$^{-1}$)** |
| Before | 6.2E+08 | 0.9 | 139 | 137 | 24 | 2.6E+07 |
| Heat stress | 5.0E+08 | 0.4 | 160 | 148 | 7 | 7.1E+07 |
| After | 1.5E+09 | 0.4 | 145 | 140 | 17 | 8.6E+07 |

C

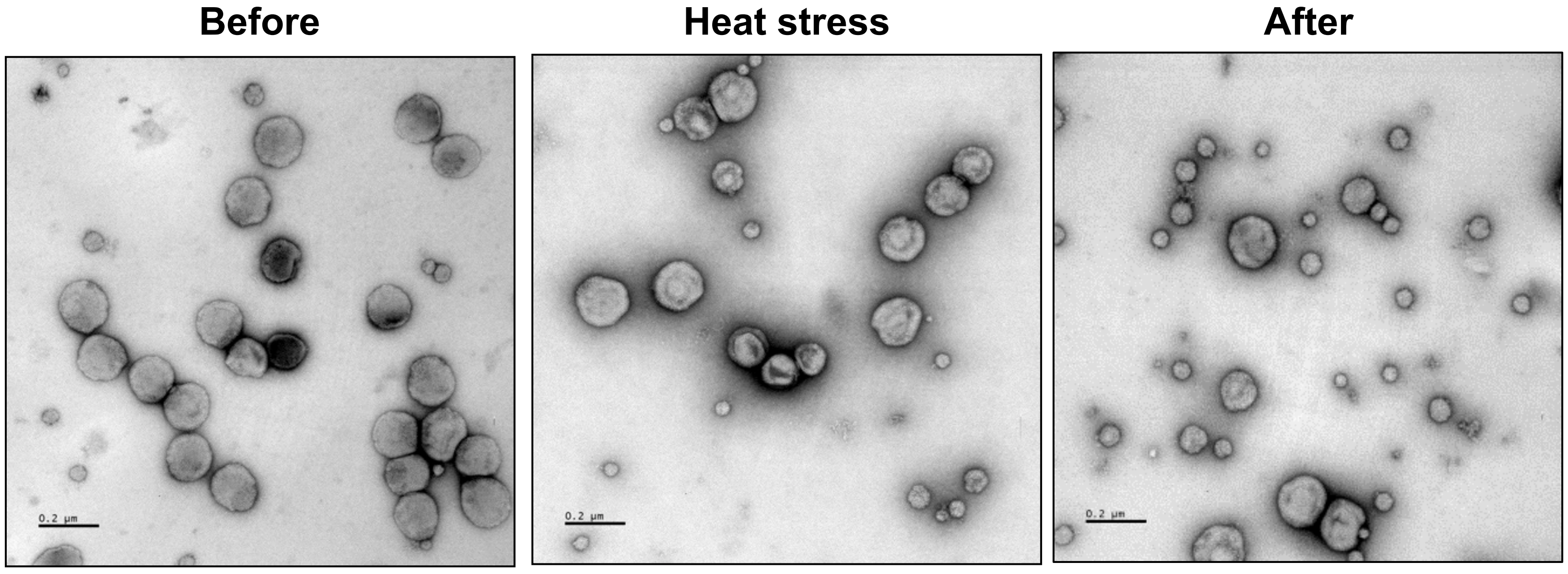


**Supplementary Figure S12. (A)** Line graph showing NTA analysis of OLN-93 EV$^{100,000\ xg}$ isolated from Before, Heat stress and After samples. A population of smaller particles was most evident in the After sample (blue box). **(B)** Table showing particle data obtained from NTA and protein concentration (BCA) assays. Note, the rate of particle accumulation in the media increased with stress. **(C)** TEM images of EV$^{100,000\ xg}$. Note the increase in number of smaller particles following stress, which supports the NTA data in (A).

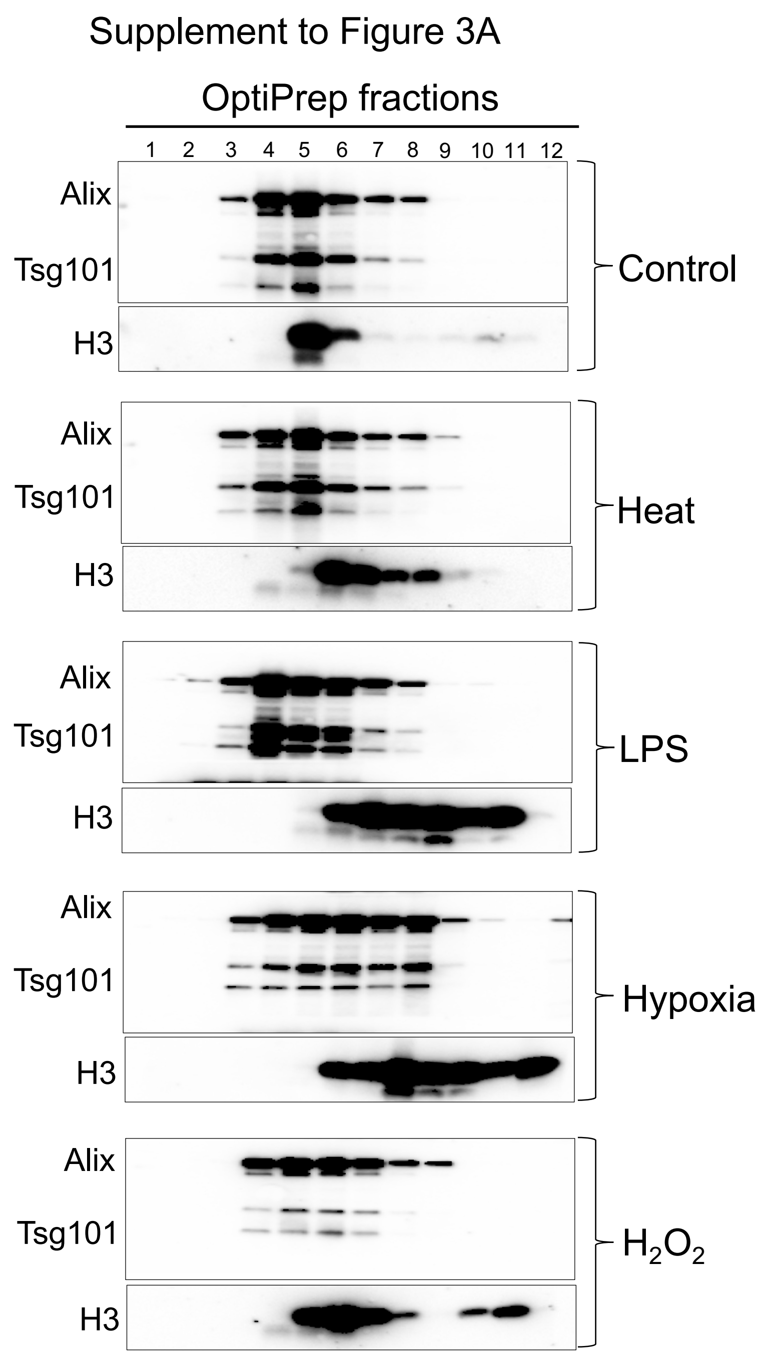


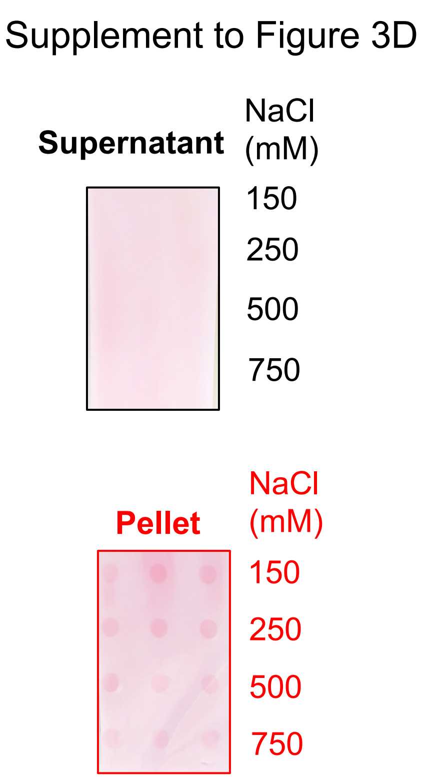


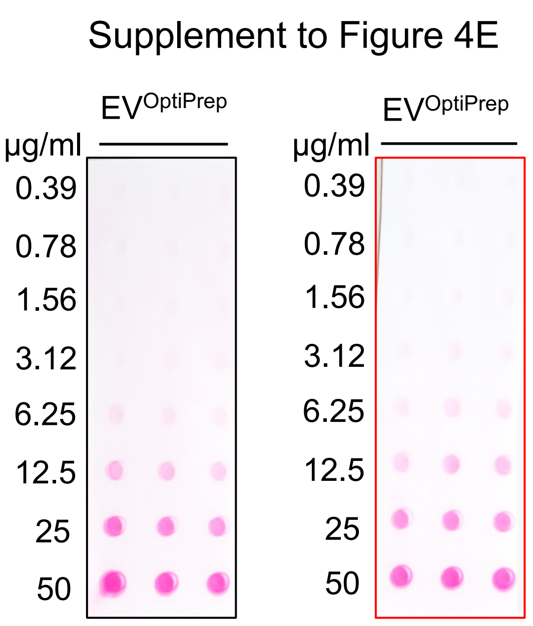


**Supplementary Figure S13.** Above (Supplement to Figure 3A), equivalent protein amount was loaded for corresponding OptiPrep fractions isolated following exposure of cells to different stress conditions (indicated on right). WB showing anti-Alix, -Tsg101 and H3. Alix and TSG staining confirm equivalent loading of EVs from different stress conditions.

Supplement to Figure 3D and 4E show Ponceau S staining of nitrocellulose membrane blots from Fig. 3D and 4E, respectively.

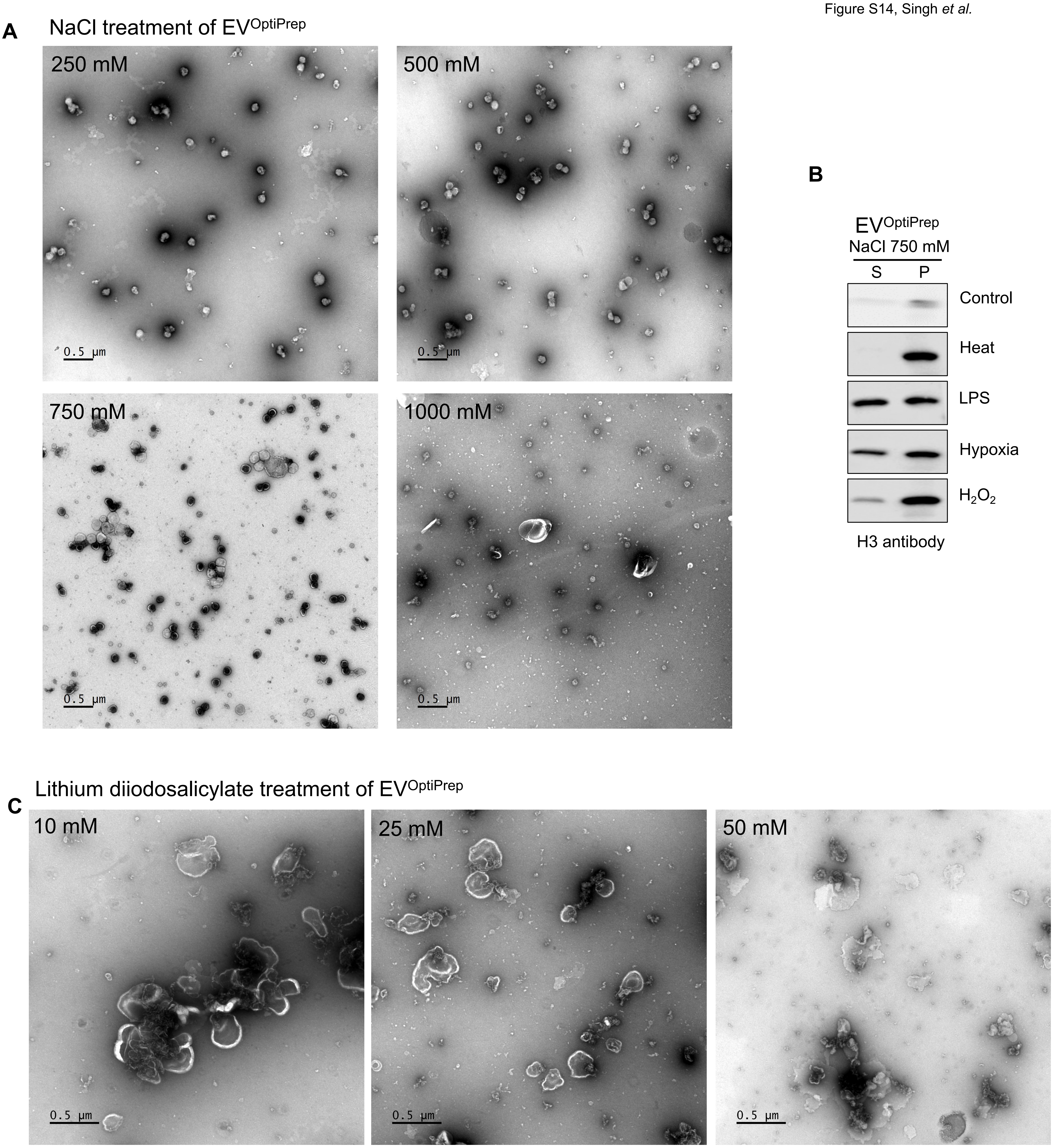


**Supplementary Figure S14. Integrity of EV$^{OptiPrep}$ treated with NaCl or lithium diiodosalicylate. (A)** NaCl treated samples were examined via TEM. **(B)** Western blot showing results of 750 mM NaCl treatment of EVs obtained following different stress conditions of OLN-93 cells. **(D)** TEM images showing integrity of EV$^{OptiPrep}$ after treatment of different lithium diiodosalicylate concentrations.

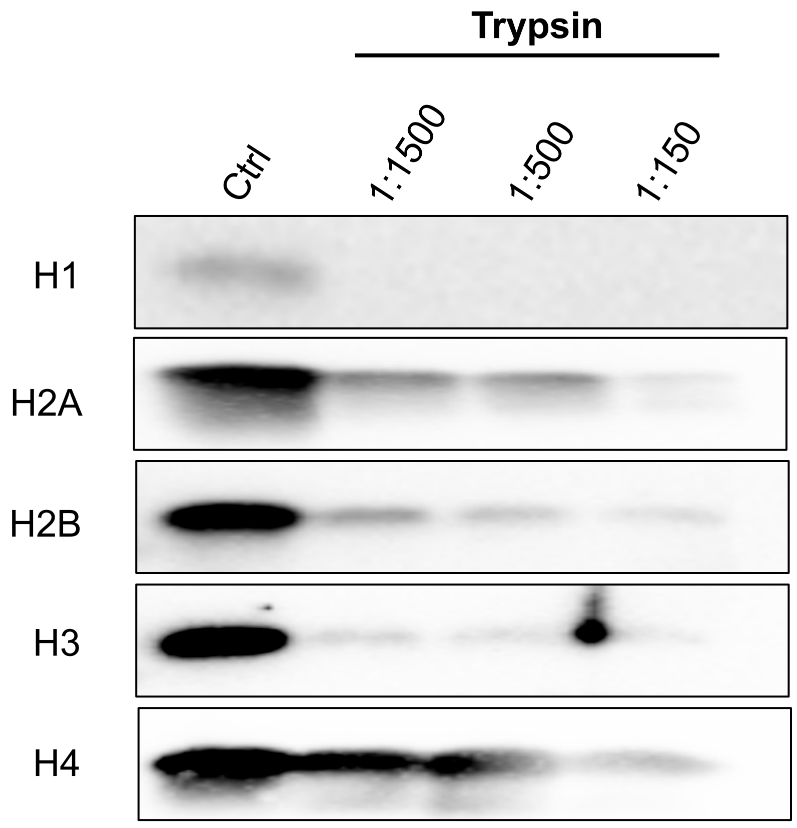


**Supplementary Figure S15.** WB showing digestion of recombinant histones using trypsin. Approximately 50 ng of each histone was treated with (1:1500-1:150 µg trypsin:µg histone). Samples were separated by SDS-PAGE and detected with the corresponding antibody to investigate pattern of digestion products. The absence of truncated digestion products suggests that free histones do not show preferential site-specific cleavage by trypsin, in contrast to EV associated histones (Fig. 3O-Q).

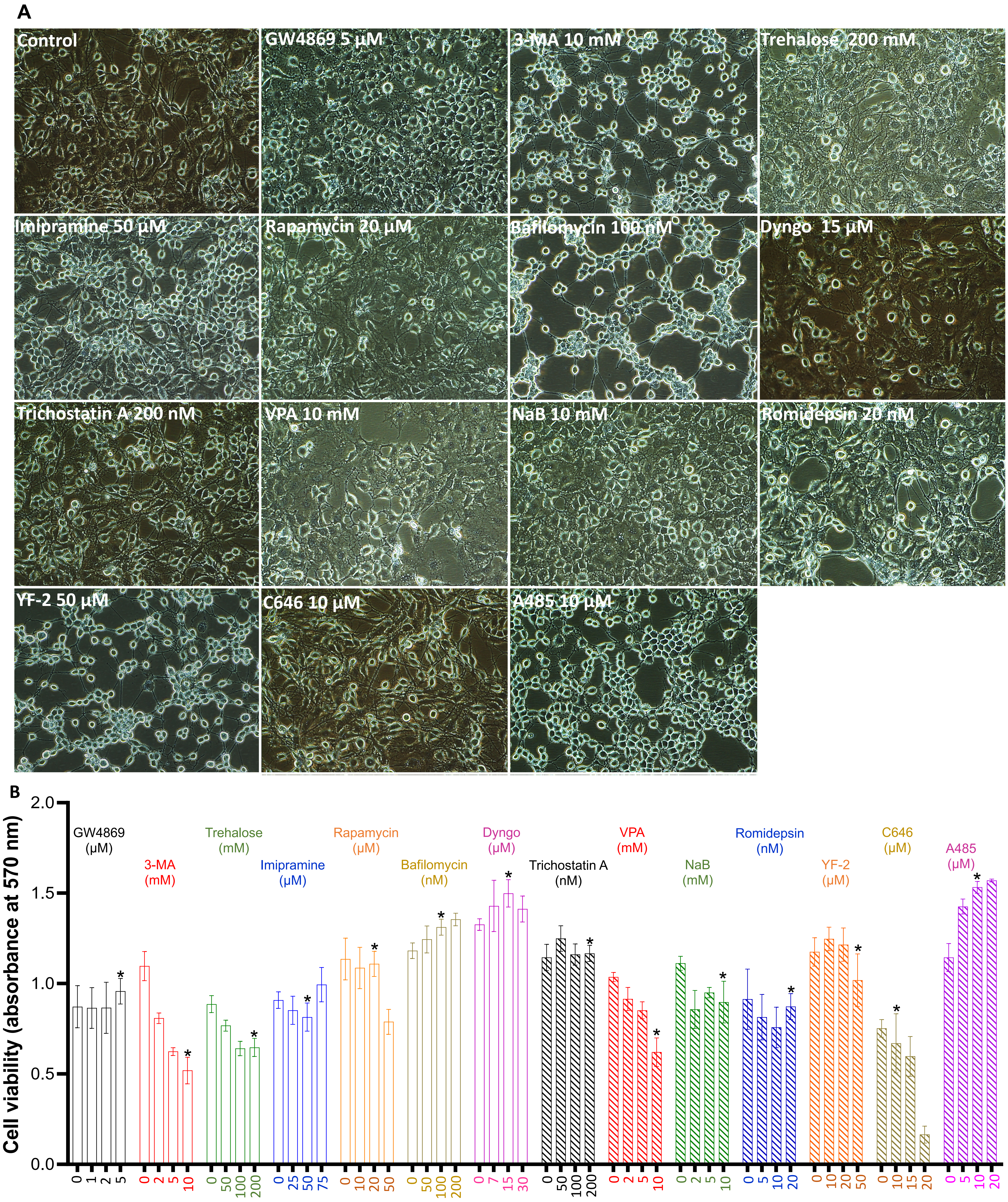


**Supplementary Figure S16. Effect of drug treatment on OLN-93 cell morphology and viability. (A)** Phase contrast images showing effect of drug treatment on cell morphology. The indicated concentration of each drug was used to analyse histone secretion in Fig. 5B. **(B)** MTT assay showing cell viability under different drug treatment conditions.

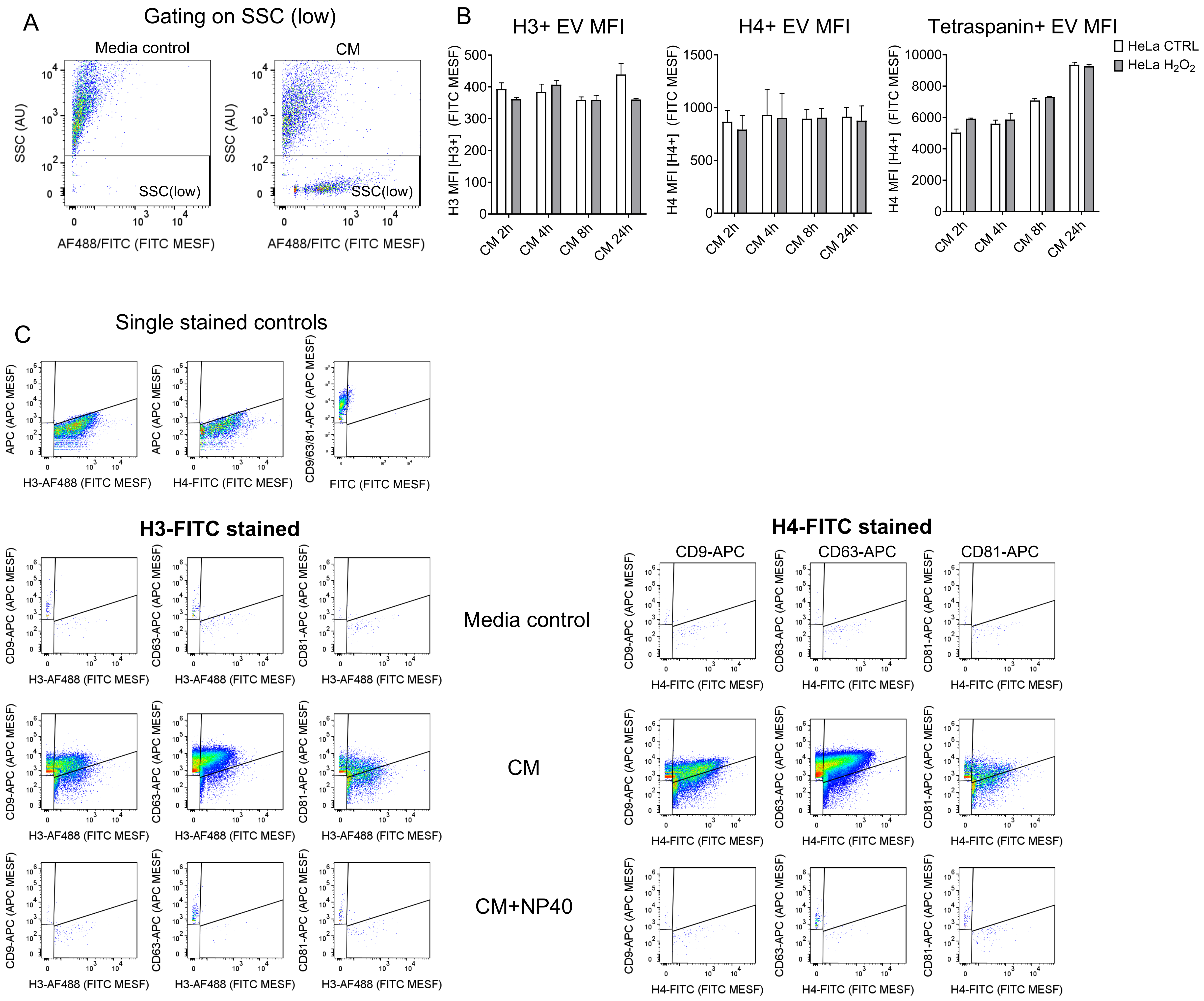


**Figure S17**: **(A)** Gating strategy used to identify SSC (low) events. **(B)** Overview of single stained controls used for gating, media controls, conditioned media (CM), and detergent controls (+NP40) for experiments using combinations of anti-histone and anti-tetraspanin antibodies, shown in Figure 6C-D.

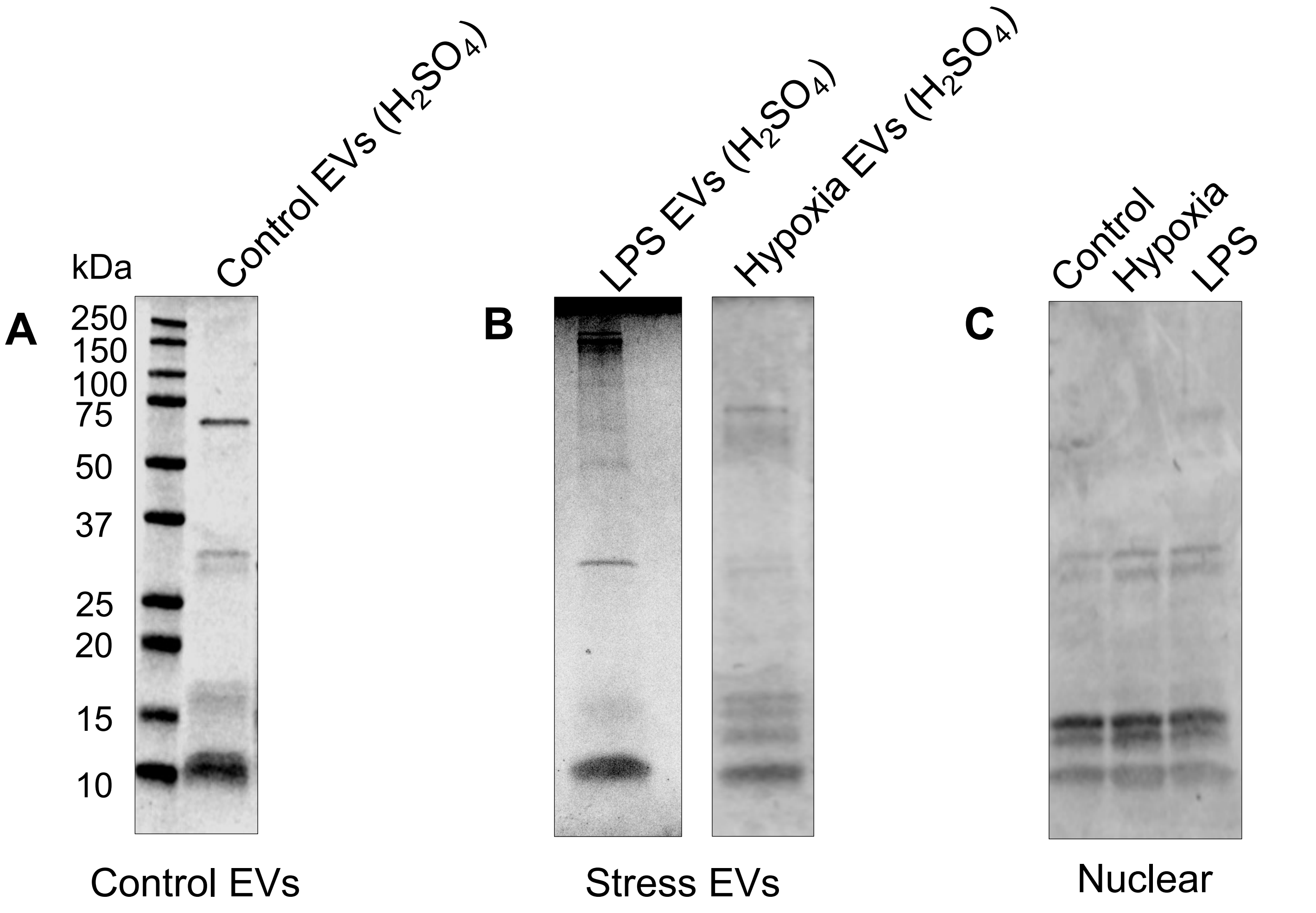


**Supplementary Figure S18. Extraction of histones from OLN-93 EV$^{OptiPrep}$ for post translational modification analyses. (A)** Control EV$^{OptiPrep}$ from OLN-93 cells (300 µg pooled from fractions 4-7) was used to acid extract histones using $H_2SO_4$. The purified proteins (1.6 µg) were separated by SDS-PAGE and stained with Comassie R-250. **(B)** As for (A) but EV histones were acid extracted from EV$^{OptiPrep}$ isolated from LPS or hypoxia conditions. **(C)** Histones acid extracted from nuclei of OLN-93 cells (as indicated).
