## Supplementary Table 1 for "Histones are exosome membrane proteins regulated by cell stress"

| Drug | Action | Concentations tested in MTT assay | Selected concentration | Reference |
|---|---|---|---|---|
| GW4869 | nSMase inhibtion. Blocks EV secretion and endocytosis. | 1,2,5 µM | 5 µM | DOI:10.1126/science.1153124 |
| 3-methyladenine | Inhibits phagophore/autophagosome formation through PI3K inhibition | 2,5,10 mM | 10 mM | DOI: 10.1073/pnas.79.6.1889 |
| D-(+)-Trehalose dihydrate | Autophagic induction (mTOR independent) | 50,100,200 mM | 200 mM | DOI: 10.1074/jbc.M609532200 |
| Imipramine | Autophagic induction (mTOR independent) / acidSMase inhibition / Inhibits macropinocytosis (less ruffling) | 25, 50, 75 µM | 50 µM | DOI: 10.1016/j.bbrc.2011.08.093 |
| Rapamycin | Autophagic induction (mTOR inhibition) | 10, 20, 50 µM | 20 µM | DOI: 10.1074/jbc.273.7.3963 |
| Bafilomycin A1 | Vacuolar ATPase inhibitor/ inhibitor of autophagosome-lysosome fusion | 50, 100, 200 nM | 100 nM | DOI: 10.1080/15548627.2015.1066957 |
| Dyngo 4a | Endocytosis inhibition through Dynamin | 7, 15, 30 µM | 15 µM | DOI: 10.1111/tra.12119 |
| Trichostatin A | HDAC class I and II inhibitor | not available | 200 nm | DOI: 10.2174%2F0929867043365099 |
| Valproic acid | HDAC class I inhibitor | 2,5, 10 mM | 10 mM | DOI: 10.1097/FPC.0b013e32835ea0b2 |
| Sodium butyrate | HDAC class I inhibitor | 2,5, 10 mM | 10 mM | DOI: 10.1016/0092-8674(78)90305-7 |
| Romidepsin | HDAC class I and II inhibitor | 5, 10, 20 nM | 20 nM | DOI: 10.1006/excr.1998.4027 |
| YF-2 | HAT inducer through CBP, PCAF, GCN5 | 10, 20, 50 µM | 50 µM | US10640457B2<br>Histone acetyltransferase activators and uses thereof |
| C646 | HAT inhibitor p300/CBP | 10, 15, 20 | 10 µM | DOI: 10.1016/j.chembiol.2010.03.006 |
| A-485 | HAT inhibitor p300/CBP | 10, 20, 40 | 10 µM | DOI: 10.1016/j.chembiol.2010.03.006 |
