## Supplementary Table 2 for "Histones are exosome membrane proteins regulated by cell stress"

**Table S2: Antibodies used in the study**

| Antibody | Company | Catalog number | Application (dilution) |
|---|---|---|---|
| Anti-histone H1 (clone AE-4,) | Millipore | 05-457 | WB (1:500) |
| anti-pan histone (clone F152.C25.WJJ) | Millipore | MABE71 | WB |
| anti-H2A acidic patch | Millipore | 07-146 | WB |
| anti-H2B | Millipore | 07-371 | WB (1:1000), ICC (1:250), TEM (1:25) |
| anti-H4 clone 62-141-13) | Millipore | 05-858 | WB (1:1000), ICC (1:250), TEM (1:25) |
| Anti-H2A | Novus | NBP1-58095 | WB (1:1000), ICC (1:250), TEM (1:25) |
| Anti-H3 DyLight 488 | Novus | NB21-1021G | FC |
| anti-H3-AF488, clone 17H2L9 | Invitrogen | MA7-02023-A488 | FC (2 ng/µL) |
| anti-H3-CT | Millipore | 07-690 | WB (1:2000), ICC (1:500), TEM (1:25) |
| Anti-H4 FITC | Novus | NB21-2044F | FC (2 ng/µL) |
| anti-H1 | Fisher Scientific | orb10806 | WB |
| Anti-H1 | Active Motif | 39708 | WB |
| Anti-H1 | Abcam | Ab154111 | TEM (1:50) |
| Anti-human CD63 (clone 12) | Fitzgerald | 435 | WB (1:1000) |
| anti-human CD63 | Thermo Fisher Scientific | MA119281 | TEM (1:50), WB (1:1000) |
| anti-rat CD63 | Novus | NBP2-68077 | WB (1:1000) |
| anti-CD63-APC clone H5C6 | Miltenyi Biotech | 130-100-182 | FC (0.6 ng/µL) |
| Anti-CD9 C-4 | Santa Cruz | sc-13118 | WB (1:1000) |
| Anti-CD9 | Sigma-Aldrich | C9993 | WB |
| Anti-CD9 | Novus | NBP2-67310 | WB (1:1000), TEM (1:25) |
| Anti-CD9 Monoclonal Antibody | Thermo Fisher Scientific | eBioKMC8 | TEM (1:25) |
| anti-CD9-APC, clone SN4 | Miltenyi Biotech | 130-128-037 | FC (0.6 ng/µl) |
| Anti-CD81 (B11) | Santa-Cruz | sc-166029 | TEM (1:50) |
| anti-CD81-APC, clone JS64 | Beckman Coulter | A87789 | FC (0.6 ng/µl) |
| anti-GFP (Clones 7.1 and 13.1, | Roche | 11814460001 | WB |
| anti-actin, beta | Sigma-Aldrich | A1978 | WB |
| anti-acetylcholine esterase (AChE) | Abcam | ab97299 | WB (1:500) |
| anti-TSG101 | Abcam | ab83 | WB (1:1000) |
| anti-GM130 | BD Transduction | 610823 | WB (1:500) |
| anti-HSP60 | Cell Signaling | 4869 | WB (1:500) |
| anti-hsp70 | Stressgen Bioreagents | SPA-812 | WB (1:500) |
| anti-annexin A1 | R&D Systems | AF3770 | WB (1:1000), ICC (1:250) |
| anti-MVP | Thermo Fisher Scientific | JM74-73 | WB (1:1000) |
| anti-Argonaute-2 | Abcam | EPR10411 | WB (1:1000) |
| anti-His tag | Cell Signaling | 2365S | WB (1:1000) |
| anti-HA tag | Cell Signaling | C29F4 | WB (1:1000) |

WB = Western blot. FC = Flow cytometry. TEM = Immuno-transmission electron microscopy, ICC = Immuno cytochemistry
